# Diverse roles of *MAX1* homologues in rice

**DOI:** 10.1101/2020.08.12.248138

**Authors:** Marek Marzec, Apriadi Situmorang, Philip B. Brewer, Agnieszka Brąszewska-Zalewska

## Abstract

Cytochrome P450 enzymes encoded by *MORE AXILLARY GROWTH1* (*MAX1*)-like genes produce most of the structural diversity of strigolactones during the final steps of strigolactone biosynthesis. The diverse copies of *MAX1* in *Oryza sativa* provide a resource to investigate why plants produce such a wide range of strigolactones. Here we performed *in silico* analyses of transcription factors and microRNAs that may regulate each rice *MAX1*, and compared the results with available data about *MAX1* expression profiles and genes co-expressed with *MAX1* genes. Data suggest that distinct mechanisms regulate the expression of each *MAX1*. Moreover, there may be novel functions for *MAX1* homologues, such as the regulation of flower development or responses to heavy metals. In addition, individual *MAX1s* could be involved in specific functions, such as the regulation of seed development or wax synthesis in rice. Our analysis reveals potential new avenues of strigolactone research that may otherwise not be obvious.

## 1. Introduction

Strigolactones (SLs) were first discovered as stimulators of seed germination in species of the genera *Orobanche, Phelipanche* and *Striga* [1]. Then, exudation of SLs from roots was found to promote hyphal branching of arbuscular mycorrhizal (AM) fungi [2]. More recently, SLs were described as a novel group of endogenous plant hormones, based on the analysis of mutants with semi-dwarf and high-branched phenotype [3,4]. Henceforth, additional roles of SLs in plant growth and development, such as the regulation of the root system development [5], elongation of the mesocotyl and stem [6,7], vasculature formation and secondary growth [8,9] and leaf senescence [10], were discovered. Finally, it was postulated that SLs are involved in plant responses to various biotic [11,12] and abiotic [13] stresses. Most SL biosynthesis appears to occur in vasculature, with resultant transport upwards in shoots or exudation out of roots [14].

More than 30 natural SLs have been identified so far, and are classified into two groups, canonical SLs and non-canonical SLs, based on their chemical structures [15] (Figure 1). All 23 so-far-identified canonical SLs contain ABC-rings connected, via an enol-ether bridge, to a methylbutenolide D-ring [16]. Conversely, a growing list of non-canonical SLs do not contain the ABCD-ring structure (Figure 1c,d) [15,17]. Non-canonical SLs were revealed for the first time when carlactone (CL) was identified [18], which was later found to be a precursor for methyl-carlactonoic acid [19] (Figure 1c). Next, zealactone [20,21] (Figure 1d) and avenaol [22] were isolated from root exudates of maize (*Zea mays*) and black oat (*Avena strigose*), respectively. These non-canonical SL lack A-, B- and C-ring but still exhibited activity during parasitic weed seed germination stimulation, which is dependent on the presence of the enol-ether-D-ring moiety in SL compounds [20,21]. This is why it is now postulated that SLs should be defined as bioactive ‘carotenoid-derived molecules with a butenolide D-ring’ [23].

**Figure 1.**
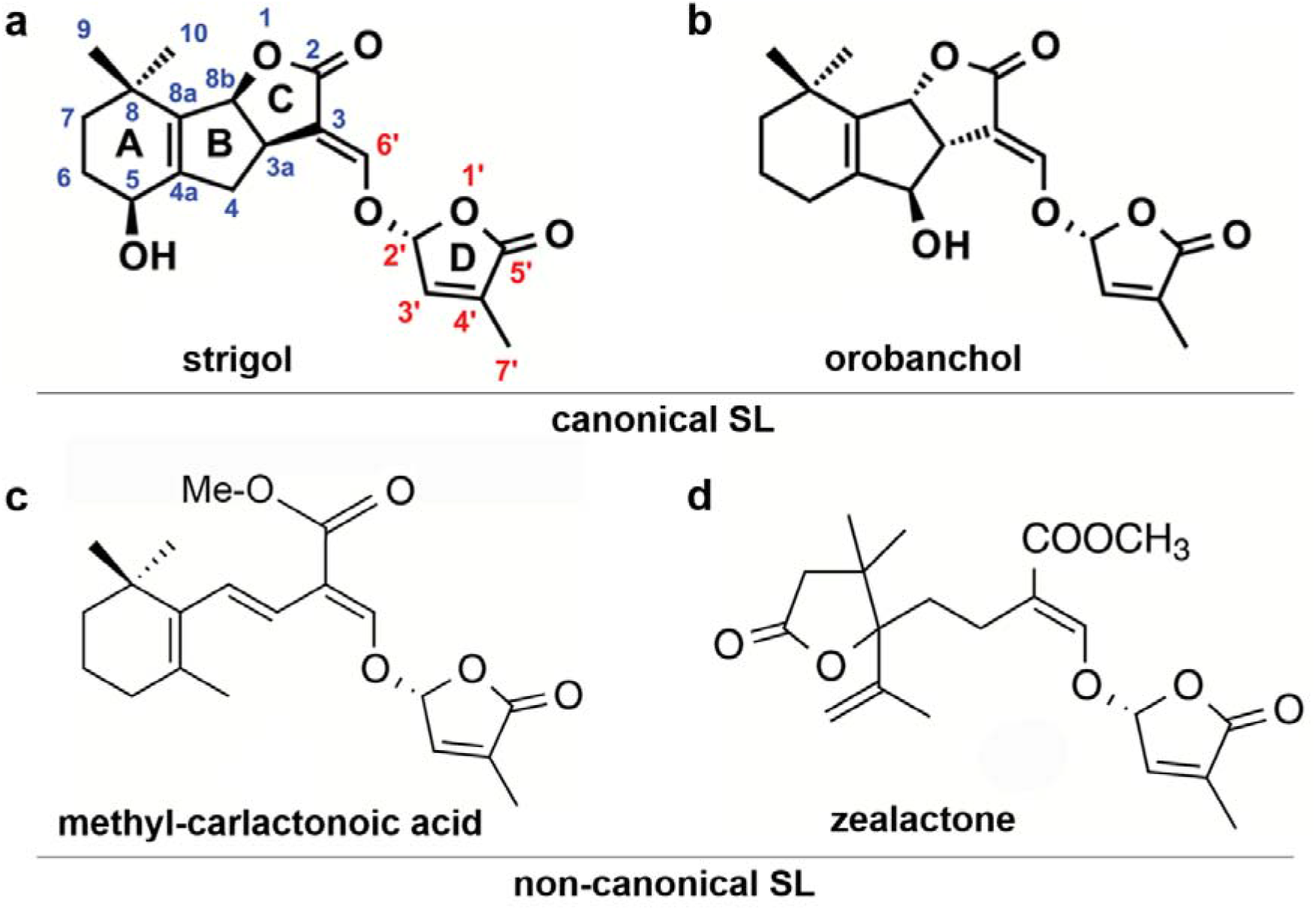
Structures of SLs. (**a, b**) canonical SLs and (**c, d**) non-canonical SLs

Canonical SL can be divided into two subgroups, strigol-type and orobanchol-type, based on the stereochemistry of the C-ring (Figure 1a,b). Strigol-type SLs have a β-oriented C-ring (3a*R*,8b*S*), and orobanchol-type SLs have an α-oriented C-ring (3a*S*,8b*R*) [23]. So far, in all examined exudates of different plants, a mixture of more than one SLs was identified [summarized by [23]]. Various SL profiles were described for different species, different varieties of the same species, and even for the same variety exposed for different growing conditions or at different developmental stages [24,25]. Some plant species, such as rice (*Oryza sativa*), do not produce strigol-type SLs, whereas the other plant species such as tobacco (*Nicotiana tabacum*) produce both, orobanchol- and strigol-type SLs [26]. Also, plants that produce both canonical and non-canonical SL were described (reviewed by [23]).

Structural diversity of SLs raises questions about the effect of the chemical structure on biological activities of SL compounds. For all the described SLs, a stimulus effect on parasitic weed seed germination, induction of hyphal branching of AM fungi and inhibition of plant shoot branching have been shown. However, different SLs exhibited various efficiency in these processes. For example, in garden pea (*Pisum sativum*), strigol and orobanchol showed less activity in inhibition of shoot branching in comparison to orobanchyl acetate and 5-deoxystrigol [27].

The large number of SLs that are produced in plants lies in contrast to the low number of components for SL biosynthesis (reviewed by [28]) and signalling (reviewed by [29,30]) pathways. The first steps of SL biosynthesis occur in plastid membranes where the D27 (DWARF27) carotenoid isomerase converts all-*trans*-β-carotene into 9-*cis*-β-carotene (Figure 2). 9-*cis*-β-Carotene is a substrate for the next stages of SL production conducted by carotenoid cleavage dioxygenases (CCDs). The first one, CCD7, is a stereo-specific dioxygenase that cleaves 9-*cis*-β-carotene to produce 9-*cis*-β-10-carotenal, which is subsequently processed by CCD8 to produce CL [18]. Genes that encode D27 and both CCDs have been identified in many different plant species, including moss (*Physcomitrella patens*). However, a different number of gene copies encoding these proteins was identified in different species. For example, two, four and six copies of *CCD8* were found in maize, rice and sorghum (*Sorghum bicolor*), respectively (reviewed by [23]). The plastidic pathway of SL biosynthesis ends with CL production and, based on grafting studies, CL and downstream products are free to move out of cells [31–33]. The structure of CL is similar to that described for canonical and non-canonical SLs, because it consists of a C19-skeleton and a C14-moiety that corresponds to the D-ring of SLs [18].

**Figure 2.**
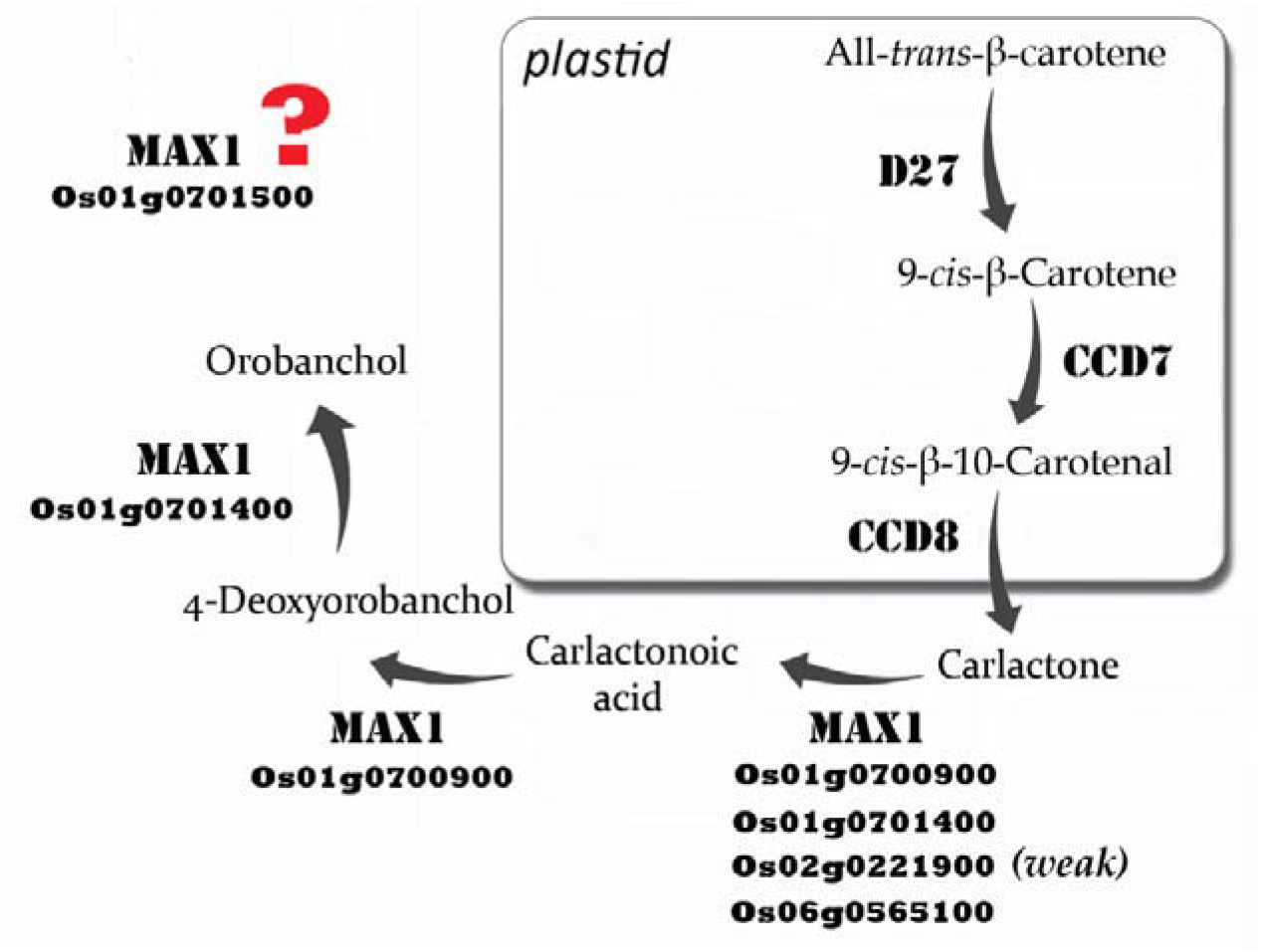
SL biosynthesis pathway in rice

The next stages of SL biosynthesis are different in the two model species, *Arabidopsis thaliana* and rice, yet many of the steps are provided by MAX1 (MORE AXILLARY GROWTH1)-like enzymes that are members of the CYP711A cytochrome P450 family. In *A. thaliana* only one copy of *MAX1* was identified and the enzyme converts CL into carlactonoic acid (9-desmethyl-9-carboxy-carlactone; CLA) [19,31]. CLA is then converted by unknown methyl transferase to methyl carlactonoate (MeCLA) [19], which is next processed into hydroxyl-methyl-carlactonoate (1’-HO-MeCLA) by a 2-oxoglutarate-dependent dioxygenase, LATERAL BRANCHING OXIDOREDUCTASE (LBO) [17,33].

The CL to CLA reaction seems to occur in all plant species so far tested [34] and CLA now appears to be the universal precursor for all SLs - both canonical and non-canonical[34]. CLA is probably not bioactive, but is converted into bioactive SLs [19]. The production of canonical SLs seems to be absent in *A. thaliana* [35]. The *max1* knockout mutant phenotypes in *A. thaliana*, rapeseed (double mutant) and tomato appear strong, suggesting that most, if not all, SL biosynthesis occurs via MAX1 [31,36,37]. CLA variants hydroylated at the A-ring have also been detected [17]. It is unclear if these CLA variants come from carotenoid precursors via the CCDs or if they are produced from CLA by other enzymes. MAX1 from *Lotus japonicus* actually converts CLA to a CLA variant hydroylated at the A-ring, at C-18 [38]. This 18-hydroxy-CLA is further converted to 5-deoxystrigol and lotuslactone by unknown enzymes. A second *L. japonicus* MAX1 has not yet been tested [38], but is a good candidate for further biosynthesis.

Five copies of *MAX1* in rice were identified, *Os01g0700900, Os01g0701400, Os01g0701500, Os02g0221900* and *Os06g0565100* (Figure 2). CL was converted into CLA by Os01g0700900, Os01g0701400, Os02g0221900 or Os06g0565100 (live yeast cell assays), but the activity of Os02g0221900 was very weak and Os01g0701500 was absent [35]. Interestingly, a maize protein closely related to Os02g0221900 also showed only very weak CL to CLA activity [35]. This Os02g0221900 sub-clade is quite distinct (Figure 3), so it will be important to discover if the members have any other distinctive function. Note that cytochrome P450s enzymatic assays require microsomes or live cells (yeast, *E. coli* or insect) (MAX1 enzymes have an N-terminal transmembrane domain) and co-expression/incubation with an NADPH-P450 reductase.

**Figure 3.**
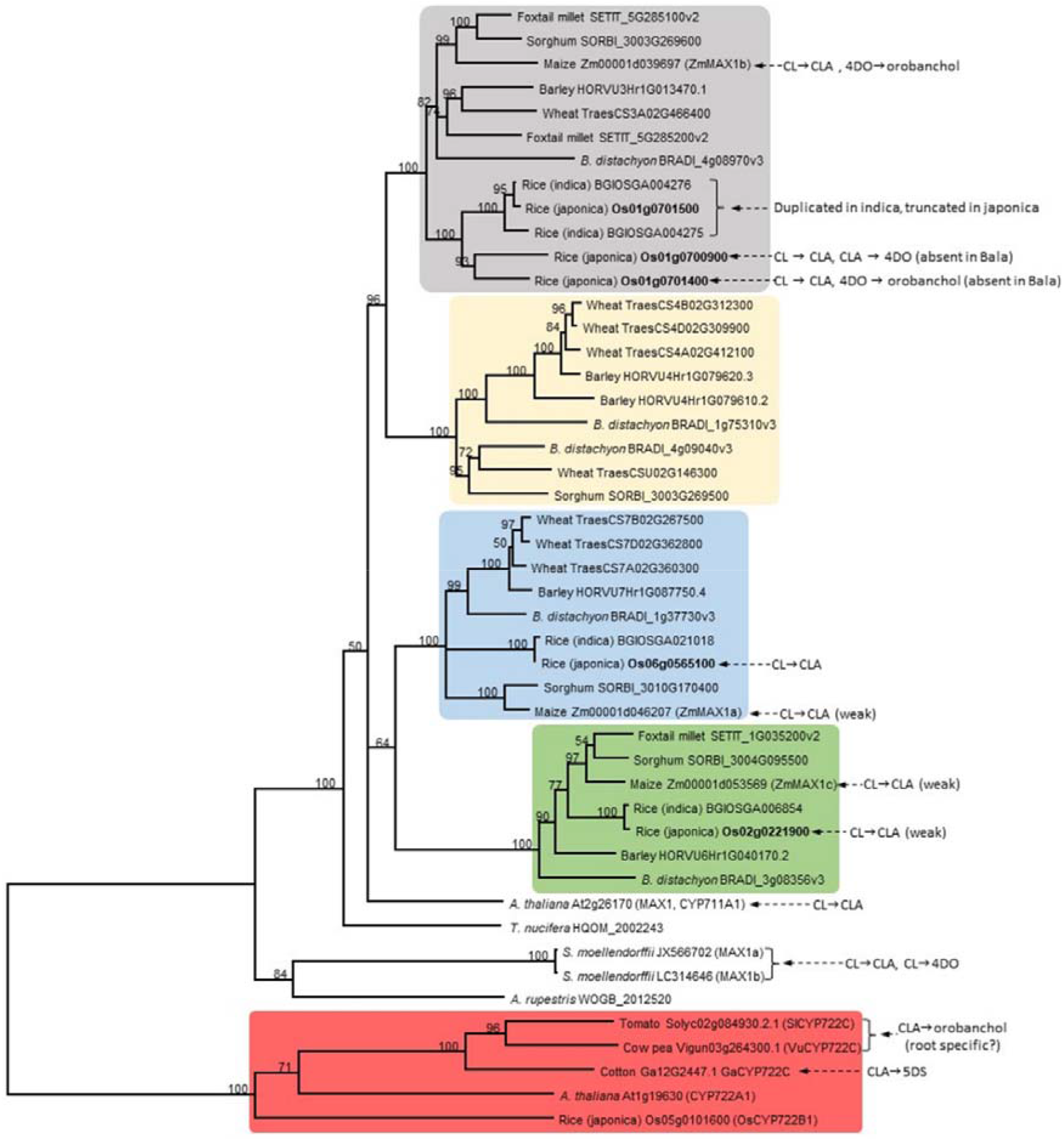
Phylogenetic relationship of MAX1 amino acid sequences from rice (*Oryza sativa*, subsp. *japonica* and *indica*), barley (*Hordeum vulgare*), wheat (*Triticum aestivum*), sorghum (*Sorghum bicolor*), maize (*Zea mays*), foxtail millet (*Setaria italica*), *Brachypodium distachyon, A. thaliana*, conifer *Torreya nucifera*, fern ally *Selaginalla moellendorffii* and moss *Andreaea rupestris*. The tree is rooted with selected CYP722C members (red) and annotated with known enzyme reactions (CL = carlactone, CLA = carlactonoic acid, 4DO = 4-deoxyorobanchol, 5DS = 5-deoxystrigol). Bootstrap percentage values shown and highlighted groups indicate distinctive clades.

Remarkably, in addition to making CLA, Os01g0700900 also independently closes the B-C rings of CLA to make 4-deoxyorobanchol (4DO; also called *ent*-2⍰-*epi*-5-deoxystrigol) [19,35]. And *Os01g0701400* can independently convert 4DO to orobanchol via hydroxylation [19,35,39]. It is unknown whether Os01g0701500, Os02g0221900 or Os06g0565100 perform steps other than CL to CLA or whether Os01g0701500 is functional (see below). The Bala indica rice cultivar (and most indica varieties) has a deletion of Os01g0700900 and *Os01g0701400* (and duplication of Os01g0701500), is high tillering, semi-dwarf and defective in SL production, yet it still produces a small amount of SLs, suggesting partial redundancy by other enzymes [40].

Non-MAX1 cytochrome P450s may also act in SL biosynthesis, and may fill this redundancy role. CYP722C from cowpea and tomato can convert CLA into orobanchol, probably via 18-hydroxy-CLA [41] (Figure 3). The knockout mutant in tomato had normal branching and normal SL feedback, but root exudates were deficient in orobanchol and solanacol, and were poor in germinating parasitic weed seeds [41]. Roots showed an increase in CLA [41], suggesting a specific function for converting CLA to orobanchol in roots. In contrast, CYP722C from cotton converts CLA into 5-deoxystrigol via 18-hydroxy-CLA [42]. Other families of enzymes, such as 2-oxoglutarate-dependent dioxygenases, that include LBO [33], have potential to act in combination with MAX1s in SL biosynthesis [36].

Complementation studies with the *A. thaliana max1-1* mutant using *MAX1* rice sequences revealed that *Os01g0700900* and *Os01g0701400* [40], and *Os02g0221900* and *Os06g0565100* [43], were able to rescue max1-1. Whereas overexpression of *Os01g0701500* in the *max1-1* background did not rescue the mutant phenotype, which might be explained by the presence of the premature stop codon 20 residues from the end of sequences [43]. Three other SLs with unknown structure were detected in rice exudates and tentatively named methoxy-5-deoxystrigol (Me-O-5-DS) isomers [40]. However, these are likely non-canonical SLs with currently unknown biosynthesis. In addition, it is still unknown whether CLA is converted to MeCLA in rice. Added CLA was converted to MeCLA in sunflower [34], but endogenous MeCLA has otherwise only been detected in *A. thaliana* and poplar [23], and the methyl transferase that makes MeCLA from CLA remains unknown. MeCLA has been proposed to be a precursor for zealactone and heliolactone [23]. It will be important to discover if MeCLA exists in rice and if it is a substrate for rice LBO.

Strigolactone variants may have evolved to not attract parasitic weeds but still attract beneficial soil microbes. Different strigolactones may also vary in their specificity for plant traits, such as branching [27]. These complex selective pressures may result in diversification of cytochrome P450s. Mutations in new gene copies may alter enzyme function or expression. In terms of the MAX1 copies that we see in rice and other well-known grass species, there appears to be diversification into four distinct MAX1 clades [35,44] (Figure 3). The role of CYP722Cs in grasses is unknown, but they may constitute a fifth clade (Figure 3). Have these clades evolved distinct and conserved enzymatic functions in grasses? There is some information from enzymatic studies in heterologous expression systems, summarized in Figure 3. Many MAX1s can catalyse CL to CLA. Some only do this weakly, such as those from the ‘blue’ group. The ‘grey’ group seems to have an emphasis on extra specific steps for making orobanchol-type SLs. However, at this stage, there are so many missing enzymatic steps for specific SLs from each grass species that it is difficult to generalize. It could equally be that each MAX1 has evolved its own function independently such that ancestral functions are not obvious or lost. Then there is the added complication about where each SL biosynthesis enzyme is expressed. For example, it will be interesting to investigate whether CYP722C[41] homologues are active in rice and have tissue-specific expression and function.

Our knowledge of the enzymes involved in the structural diversification of SLs remains scarce. Because different SLs have different biological activities, it will be crucial to identify the components involved in the synthesis of each SL and find out if they have tissue-specific regulation. This will allow us to investigate the role of SLs in plants with much higher precision. With that in mind, we performed *in silico* analyses using all five rice *MAX1* homologues to uncover regulatory mechanisms that may control their expression. Together with the analysis of their expression patterns during plant development and responses to various factors, it is possible to predict further roles of rice *MAX1* homologues. These results might help us better understand the structural diversification of SLs and better elucidate the regulation of each enzymatic step.

## 2. Materials and Methods

### 2.1. Rice MAX1 sequences and conserved domain identification

Genomic, coding and protein sequences of rice MAX1 homologues were obtained from NCBI database (www.ncbi.nlm.nih.gov): Os01g0700900 – gene ID 4326926; Os01g0701400 – gene ID 9269315; Os01g0701500 – gene ID 4326929; Os02g0221900 – gene ID 4328761; Os06g0565100 – gene ID 4341325.

Sequence of the promotor region of rice genes were obtained using PlantPAN 3.0 platform (http://plantpan.itps.ncku.edu.tw). For each of gene the sequence of 2000 nucleotide (nt) upstream transcription start site and 500 nt downstream transcription start site were obtained.

The amino acid sequence of AtMAX1 was used as a query for BLAST search on Ensembl Plants (https://plants.ensembl.org/Multi/Tools/Blast) to gather sequences for the phylogenetic tree (Figure 3). Genes with less than 400 amino acids were excluded. MAX1 homologues from maize and *Selaginella moellendorffii* are the same as previously reported [35]. Amino acid sequences of MAX1 homologs of *Andreaea rupestris* and *Torreya nucifera* were obtained from 1KP projects https://db.cngb.org/onekp/ [44]. Amino acid sequences for CYP772 clade are the same as previously reported [41,42] [refs]. MAFFT was used for multiple sequence alignment [45]. The phylogenetic tree was constructed using Geneious Tree Builder (Geneious 8.1.9) following the Neighbour-joining method with bootstrap resampling of 1000 replicates and rooted to CYP772 clade. Sequences, alignment and gene IDs are available in Supplementary Data.

### 2.2. Searching of transcription factor motifs in promoter region of rice MAX1 genes

To predict the sites recognized by transcription factors (TFs), 2500 nt of promoter region of all five rice *MAX1* genes was used as a query in tool ‘Promoter Analysis’ implemented in PlantPAN 3.0 platform (http://plantpan.itps.ncku.edu.tw). In each case, only the database for rice-specific TFs was selected and no user-customized motifs were used. The function of TFs that are specific only to a single *MAX1* homologue, was identified based on available literature and TF databases, such as: PlantTFDB (http://planttfdb.cbi.pku.edu.cn), CIS-BP Database (http://cisbp.ccbr.utoronto.ca), New PLACE (https://www.dna.affrc.go.jp/PLACE/) and JASPAR (http://jaspar.genereg.net).

### 2.3. Identification of miRNA that regulate MAX1 genes

For identification of miRNA that may regulate the *MAX1* genes, the psRNATarget web server was used (http://plantgrn.noble.org/psRNATarget, 2017 release) [46]. The coding sequence of each *MAX1* gene was used as a query and only rice specific miRNA was searched for, according to these rules: penalty for G:U pair: 0.5; penalty for other mismatches: 1; extra weight in seed region: 1.5; seed region: 2-13nt; # of mismatches allowed in seed region: 2; HSP size: 19; penalty for opening gap: 2; penalty for extending gap: 0.5. Only results with a final score (expectation value) up to 5 were considered for further investigation. Expectation value is the penalty for the mismatches between mature small RNA and the target sequence. A higher value indicates less similarity (and possibility) between small RNA and the target candidate. The function of miRNAs that are specific only to a single *MAX1* homologue was identified based on the available literature.

### 2.4. Profile expression of MAX1 genes

Data of *MAX1* genes expression were obtained from the Rice Expression Database (http://expression.ic4r.org) that integrates expression profiles derived entirely from NGS RNA-Seq data of rice (Nipponbare variety). The following Loc_IDs: LOC_Os01g50520, LOC_Os01g50580, LOC_Os01g50590, LOC_Os02g12890 and LOC_Os06g36920 were used for *Os01g0700900, Os01g0701400, Os01g0701500, Os02g0221900, Os06g0565100*, respectively. Only experiments containing data for all of the five *MAX1* homologues were used for *MAX1* profile expression comparisons.

### 2.4. Gene co-expression network of rice MAX1 homologues

Lists of genes co-expressed with *MAX1* homologues were obtained using RiceFREND (https://ricefrend.dna.affrc.go.jp). The locus ID of each gene was used for the ‘single guide gene’ searching option. Only genes with a Mutual Rank (MR) lower than 100 were taken for further investigation. MR is calculated as the geometric mean of the correlation rank of gene A to gene B and of gene B to gene A, based on 24 datasets representing 815 microarray data including redundant data among the datasets. Details on each dataset can be accessed from the RiceXPro database (https://ricexpro.dna.affrc.go.jp). The data have also been deposited in NCBI’s Gene Expression Omnibus (GEO) and are accessible through GEO series accession numbers, GSE21396, GSE21397, GSE36040, GSE36042, GSE36043, GSE36044, GSE39423, GSE39424, GSE39425, GSE39426, GSE39427, GSE39429 and GSE39432. Function of selected genes was described based on available literature.

## 3. Results

Protein sequences of rice MAX1 homologues exhibited a range of identity from 80.8 to 48%, and in all sequences, the conserved domain for cytochrome P450 (pfam00067) was present (Figure S1). We collected MAX1s amino acid sequences from species closely related to rice and produced a new phylogenetic tree specific for grasses (Figure 3). Interestingly, four separate MAX1 clades are obvious in the grass species. We also examined available enzymatic data and included those on Figure 3, which shows much flexibility in enzyme function. Surprisingly, the recently discovered CYP722C SL enzymes are only distantly related to MAX1s (Figure 3). Previously it was reported that Os01g0701500 has no enzymatic activity, because this sequence was unable to rescue phenotype of *max1-1* mutant in *A. thaliana* [43] and this protein did not exhibit enzymatic activities described for other MAX1s [35]. It was postulated that a lack of enzymatic activity of Os01g0701500 is due to premature stop codon, in comparison to the other MAX1 protein (Figure S1). On the other hand still, the full sequence of the conserved domain for cytochrome P450 in present in Os01g0701500 (Figure S1). This is why Os01g0701500 was included in presented studies.

### 3.1. Transcription factor motifs that are present in the promoter region of rice MAX1 genes

The promoter region of all five rice *MAX1* genes (*Os01g0700900, Os01g0701400, Os01g0701500, Os02g0221900, Os06g0565100*) were screened to identify sequences recognized by transcription factors (TFs). In each case, sequence of 2000 nucleotide (nt) upstream transcription start site and 500 nt downstream transcription start site was used. Various number of motifs recognized by TFs were identified for all genes (Table S1A-E). The largest number of putative TF binding sites was found in the promoter region of *Os06g0565100* (1309 motifs), whereas the smallest number was observed in case of *Os02g0221900* (1023). When the numbers of TFs that may bind to the identified motifs were estimated, similar values in the range from 219 to 250 were obtained for all analysed promoters (Figure 4a). Among them, four, five, 14, eight and 31 TFs were specific only to the promoter sequence of *Os01g0700900, Os01g0701400, Os01g0701500, Os02g0221900, Os06g0565100*, respectively (Figure 4b). TFs common for all *MAX1* homologues were identified (60) (Table S2), as well as TFs common for different pairs of *MAX1* (Figure 1b). The biggest number of common TF binding sites was observed in the promoter regions of *Os02g0221900* and *Os06g0565100* (10) and *Os01g0700900* and *Os01g0701400* (8), whereas the lowest similarity in the TF binding sites was present between promoter regions of *Os01g0701500* and *Os02g0221900* (1) (Figure 4b).

**Figure 4.**
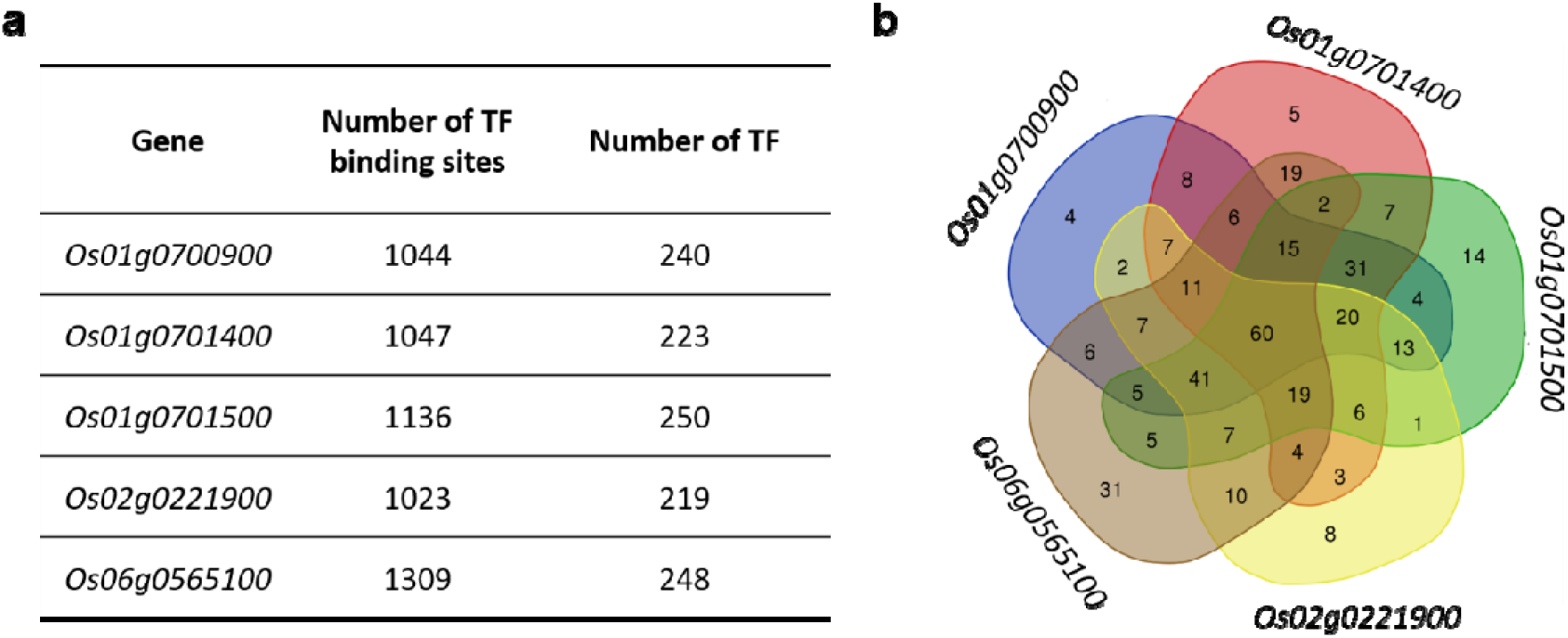
Motifs recognized by TFs that are present in the promoter region of rice *MAX1* genes. (**a**) Number of TF binding sites and TFs that bind to motifs identified in promoter region of analyzed genes. (**b**) The Venn diagram illustrates TFs that are specific to each of rice *MAX1* genes and those that are common for different *MAX1* homologues.

### 3.2. Transcription factors specific to Os01g0700900

In the case of the promoter region of *Os01g0700900*, five motifs recognized by four different TFs were identified that are specific only to this one *MAX1* gene (Table 1; Table S3). For all of them the role in plant response to cold was previously described. I.e., expression of *Os03g0820400* (TFmatrixID_0216) was induced by cold more than 4-fold in rice cold tolerant cultivar Oro (da Maia) [47]; *Os10g0377300* (TFmatrixID_0298) was typed as candidate genes related to low temperature tolerance [48]; whereas *Os12g0123700* (TFmatrixID_0381) and *Os02g0170300* (TFmatrixID_0529) were found to be cold-responsive in rice and *Oryza ofcinalis* [49]. Among other abiotic stresses, binding sites recognized by TFs involved in plant responses to drought [50–53], salt [54–56], arsenic [57], cadmium [58], nitrogen [59,60] or iron [61] deficiency were identified (Table 1; Table S3). Binding sites of TFs that play a role in plant response to pathogen attack, such as viruses [62], bacteria [63] or fungi [64], were also present. Finally, some of the TFs, that are specific only to *Os01g0700900* and are involved in developmental processes, such as root [65] or flower development [66,67], leaf senescence [68,69] and seed dormancy/germination [70] were found. Recently the function of *OsMADs57*, that exclusively bind the motif only in the promoter region of *Os01g0700900*, was described [71]. Knockdown of *OsMADs57* resulted in plant height reduction, inhibition of internode elongation and reduction in panicle exertion. Additionally, mutated plants contained less bioactive forms of gibberelins (GAs), when compared to wild-type, and were more sensitive to GA_3_ treatment [71]. This feature indicates the possible crosstalk between SL and GA biosynthesis pathways.

**Table 1.** Number of TFs found in promoter regions of *MAX1* homologues, categorised according to functions in plant development, growth and response to stresses.

|  | *Os*<br>*01g0700900* | *Os*<br>*01g0701400* | *Os*<br>*01g0701500* | *Os*<br>*02g0221900* | *Os*<br>*06g0565100* |
| --- | --- | --- | --- | --- | --- |
| ***abiotic stresses*** |  |  |  |  |  |
| iron status | 1 | 0 | 1 | 0 | 4 |
| nitrogen status | 2 | 0 | 2 | 0 | 11 |
| phosphorus status | 0 | 1 | 0 | 0 | 2 |
| arsenic | 1 | 1 | 0 | 0 | 9 |
| cadmium | 2 | 0 | 1 | 0 | 8 |
| chromium | 0 | 0 | 1 | 1 | 0 |
| cold | 4 | 2 | 1 | 0 | 7 |
| drought | 2 | 1 | 3 | 1 | 20 |
| salt | 2 | 1 | 3 | 0 | 7 |
| submergence | 0 | 1 | 1 | 0 | 3 |
| ***biotic stresses*** |  |  |  |  |  |
| bacteria | 1 | 1 | 1 | 0 | 11 |
| viruses | 1 | 0 | 2 | 0 | 0 |
| fungi | 1 | 0 | 2 | 0 | 17 |
| insects | 0 | 0 | 0 | 0 | <u>11</u> |
| ***developmental processes*** |  |  |  |  |  |
| plant height | <u>1</u> | 0 | 0 | 0 | 0 |
| shoot architecture | 0 | 0 | 0 | 0 | <u>6</u> |
| root development | 1 | 1 | 2 | 1 | 6 |
| flower development | 1 | 1 | 3 | 2 | 2 |
| seed development | 0 | 0 | 0 | <u>2</u> | 0 |
| seed dormancy | 1 | 0 | 0 | 1 | 0 |
| leaf senescence | 1 | 0 | 1 | 0 | 0 |
| secondary wall formation | 0 | 0 | <u>4</u> | 0 | 0 |
| wax synthesis | 0 | 0 | 0 | 0 | <u>1</u> |
| ***hormonal cross talk*** |  |  |  |  |  |
| abscisic acid | 0 | 1 | 0 | 0 | 1 |
| ethylen | 0 | 1 | 0 | 0 | 1 |
| gibberelins | 1 | 0 | 0 | 0 | 1 |

Bolded and underlined text indicates that this function was specific only to one *MAX1* homolog.

### 3.3. Transcription factors specific to Os01g0701400

Analysis of the *Os01g0701400* promoter region revealed five TF motifs that were not present in other *MAX1* homologues (Table 1; Table S4). TFs that belong to the TFmatrixID_0503 group were identified as regulators of flowering [67,72–76]. I.e., *Os02g0682200* (*OsMADS6*) and *Os04g0580700* (*OsMADS17*) that are involved in the specification of floral organ identity, whereas *Os08g0531700* (*OsMADS7*) and *Os09g0507200* (*OsMADS8*) are involved in flower development [67]. Among other developmental processes, some of identified TFs play a role in shaping of root architecture [77,78] (Table S4). A wide range of TFs that may bind the promoter of *Os01g0701400* were identified as related to abiotic stresses, including response to arsenic [57,79], cold [80], drought [51–53,81–83], phosphate deficiency [84] and submergence [85]. On the other hand, four genes that belong to the AP2 family (Os03g0183200, Os07g0617000, Os09g0286600 and Os09g0287000), were previously described as induced by infection of *Xanthomonas oryzae* pv. *oryzae* [63]. Additionally two motifs, with unknown TFs were found: GLUTEBP2OS and ANAERO5CONSENSUS (Table S4). The first one regulates the transcription of genes encoding glutelin storage proteins [86], whereas the latter one was found *in silico* in promoters of 13 anaerobic genes involved in the fermentative pathway [87]. Finally, two TFs that may link SL biosynthesis with other phytohormones were identified. *OsERF67* (*Os07g0674800*) was significantly increased by ethylene and abscisic acid treatment, whereas treatment with auxin, GA or brassinosteroid did not affect the expression of *OsERF67*.

### 3.4. Transcription factors specific to Os01g0701500

In the promoter region of *Os01g0701500* the second highest number of specific TF binding motifs were found, however some TF groups (TFmatrixID_0386, TFmatrixID_0388, TFmatrixID_0389, TFmatrixID_0390, TFmatrixID_0395; as well as TFmatrixID_0524, TFmatrixID_0547) were represented by the same representatives (Table S5). Identified TFs, were involved in plant responses to various biotic stresses, including viruses [62], bacteria [88], and fungi [64], and different abiotic stresses (Table S5). Among TFs related to abiotic stresses, those involved in iron [61,89] and nitrogen deficiency [59,60,90], response to cold [91], drought [50,51,92,93], salt [54,56,94], cadmium [58], chromium [95], submergence [96] were found. Binding motifs for large number of TFs involved in developmental processes, including flower development [72,97–102], root development [77] and leaf senescence [103,104] (Table 1; Table S5) were identified in the promoter region of *Os01g0701500*. Additionally binding sites for TFs that are involved in secondary wall formation were identified [105,106] (Table S5). Two of them, *Os01g0701500* and *Os06g0131700*, are known as OsSWN1 and OsSWN2, respectively (SECONDARY WALL NAC DOMAIN PROTEIN1/2). It was shown that overexpression of OsSWN1 strongly induced ectopic secondary wall formation [107]. Two other TFs, *Os06g0104200* and *Os08g0103900* were able to bind the promoter region of AtMYB46 and functionally complement the *A. thaliana snd1/nst1* double mutant that exhibits lack of lignified secondary walls in fibres. Overexpression of *Os06g0104200* and *Os08g0103900* in *A. thaliana* causes ectopic deposition of cellulose, xylan and lignin [108]. Those results indicate that one of the *MAX1* homologues - *Os01g0701500* may be regulated by the group of TFs related to secondary wall formation.

### 3.5. Transcription factors specific to Os02g0221900

In the promoter region of *Os02g0221900*, motifs recognised by five different TFs, that are specific only for this one *MAX1* gene, were identified (Table 1; Table S6). Interestingly none of the TFs that were involved in plant response to biotic stresses had been found. In the aspect of TFs related to abiotic stresses, those involved in response to cold [109], drought [110] and chromium [95] were identified. Whereas among TFs related to developmental processes, those involved in root development [77,111], flower development [112,113] were able to bind promoter of *Os02g0221900*. Also the motif that is recognized by *OsVP1* (*Os01g0911700*), rice orthologue of Arabidopsis ABI3, is present in the promoter of *Os02g0221900*. Experimental data indicate that *OsVP1* is a major determinant of seed specificity and regulate the spatial pattern of expression of genes in developing seed [114]. Additionally only this one MAX1 homolog is under control of *OsTCP5* (*Os01g0763200*), a TF which belongs to the cell division-regulating TCP family, that is regulated by SL and CK [6]. It was proved that *OsTCP5* is involved in SL-controlled axillary bud outgrowths [115] and in the control of cell division in rice mesocotyl, that depends on SL and CK [6].

### 3.6. Transcription factors specific to Os06g0565100

The highest number of TF binding sites were found in the promoter region of *Os06g0565100* (1309) (Table S1) and also the highest number of TFs specific only for this MAX1 homolog was identified (Table S7). Similar to the previous genes that were analysed, *Os06g0565100* may be also under control of TFs involved in plant response to bacteria [116,117], fungi [63,64,118–120] and insects [121,122], as well as cold [48,49,80,80,123,124], drought [50–53,82,83,110], cadmium [58,125], submergence [87,126,127], salt [81,128], arsenic [57,79], or iron [129], nitrogen [59,60,90,130] and phosphorus [84] deficiency. In the promoter region of *Os06g0565100* motifs recognized by TFs are related to flower [112,131] and root [77,111,132] development, or controlling shoot architecture [133]. Among TFs that are able to bind promoter of *Os06g0565100*, four ethylene response TFs were found: Os02g0202000 (OsWR1), Os06g0604000 (OsWR2), Os02g0797100 (OsWR3) and Os06g0181700 (OsWR4). Functional analysis of OsWR1 indicated that this TF is a positive regulator of drought resistance in rice, because it promotes expression of genes that are involved in wax synthesis, and therefore involved in water loss reduction [134]. Additionally it was shown that transcript levels of *OsWR1* were induced by drought, salt and ABA treatment [134]. On the other hand OsAP2-39 (Os04g0610400), that may recognise ten motifs in the promoter region of *Os06g0565100*, was described as a key regulator of the interaction between ABA and GAs in rice [135]. Overexpression of *OsAP2-39* results in yield reduction by decreasing the number of seeds and upregulation of ABA biosynthesis *via* increased activity of *OsNCED-I* gene (encoding 9-cis-epoxycarotenoid dioxygenase) involved in this process. Finally, overexpression of *OsAP2-39* upregulates the expression of the ELONGATION OF UPPER MOST INTERNODE I (EUI) gene that is involved in the epoxidation of the active GAs, and thus reduces the level of bioactive GAs in rice [135].

### 3.7. MiRNAs that may bind rice MAX1 homologues

The highest number of miRNA targets was identified for *Os01g0700900* – 12, for the other *MAX1s* the number of miRNA target sites ranged from eight to 11 (Figure 5a, Table S8). In the case of miRNA targets identified for *Os01g0700900, Os01g0701400* and *Os01g0701500*, each target site was recognized by various miRNAs. In the sequence of *Os02g0221900* four target sites for osa-miR5075 were found, whereas for in the sequence of *Os06g05651000*, two targets for osa-miR2927, osa-miR5075, osa-miR3980a-3p and osa-miR3980b-3p were identified (Table S8). Among all identified miRNAs, eight were specific only to *Os01g0700900*, six for *Os01g0701500*, five for *Os02g0221900*, and four for *Os01g0701400* and *Os06g05651000* genes (Figure 5b). None of the miRNAs identified match four or all five rice *MAX1* genes. Only two miRNA match motifs in three *MAX1* homologues: osa-miR419 (*Os01g0700900, Os01g0701400, Os01g0701500*) and osa-miR5075 (*Os01g0701500, Os02g0221900, Os06g0565100*) (Table S8).

**Figure 5.**
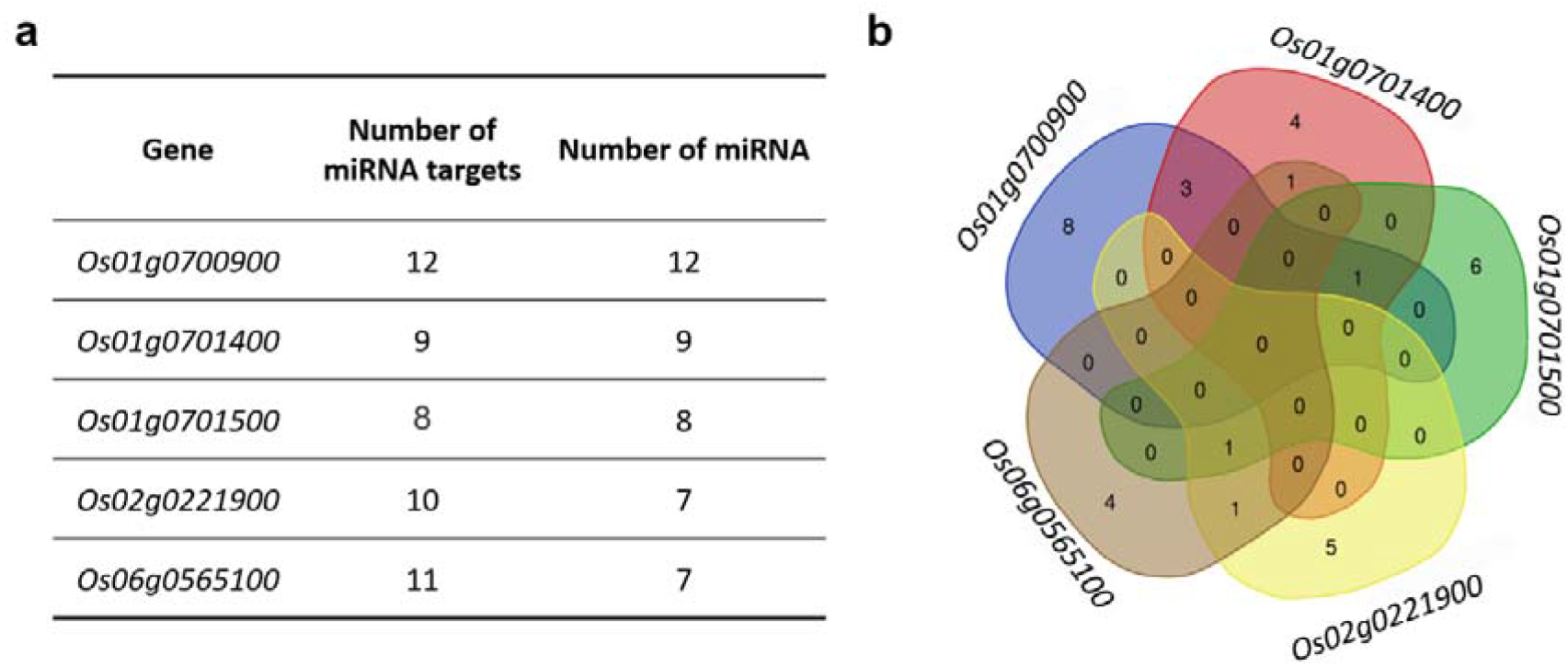
MiRNAs which recognize targets that are present in sequence of rice MAX1 genes. (**a**) Number of miRNA targets found in MAX1 homologs and number of miRNA that target sequences of MAX1s. (**b**) The Venn diagram illustrates miRNA that are specific to each of rice MAX1 genes and those that are common for different MAX1 homologues.

### 3.8. MiRNA specific to Os01g0700900

Within the miRNAs identified for *Os01g0700900* only eight were specific to this one MAX1 gene. Within these miRNAs, osa-miR2055, which is expressed preferentially in rice roots, plays the role in plant response to increased temperature (Table 2). When two rice varieties were compared: tolerant to heat (Nagina 22) and susceptible (Vandana), upregulation of osa-miR2055 was observed in the heat tolerant variety in response to short (42°C/36°C day/night for 24 h) and long (42°C/36°C day/night for 5 days) heat treatment [136]. A second miRNA, specific for *Os01g0700900* was osa-miR1432-3p (Table S8), which is known to be involved in rice response to rice stripe virus (Table 2), because targets of that miRNA are well characterized disease resistance genes (*LOC_Os02g42160* and *LOC_Os07g35680*) encoding wall-associated receptor kinase-like 1 and cysteine-rich receptor-like protein kinase 8, respectively [137].

**Table 2.**
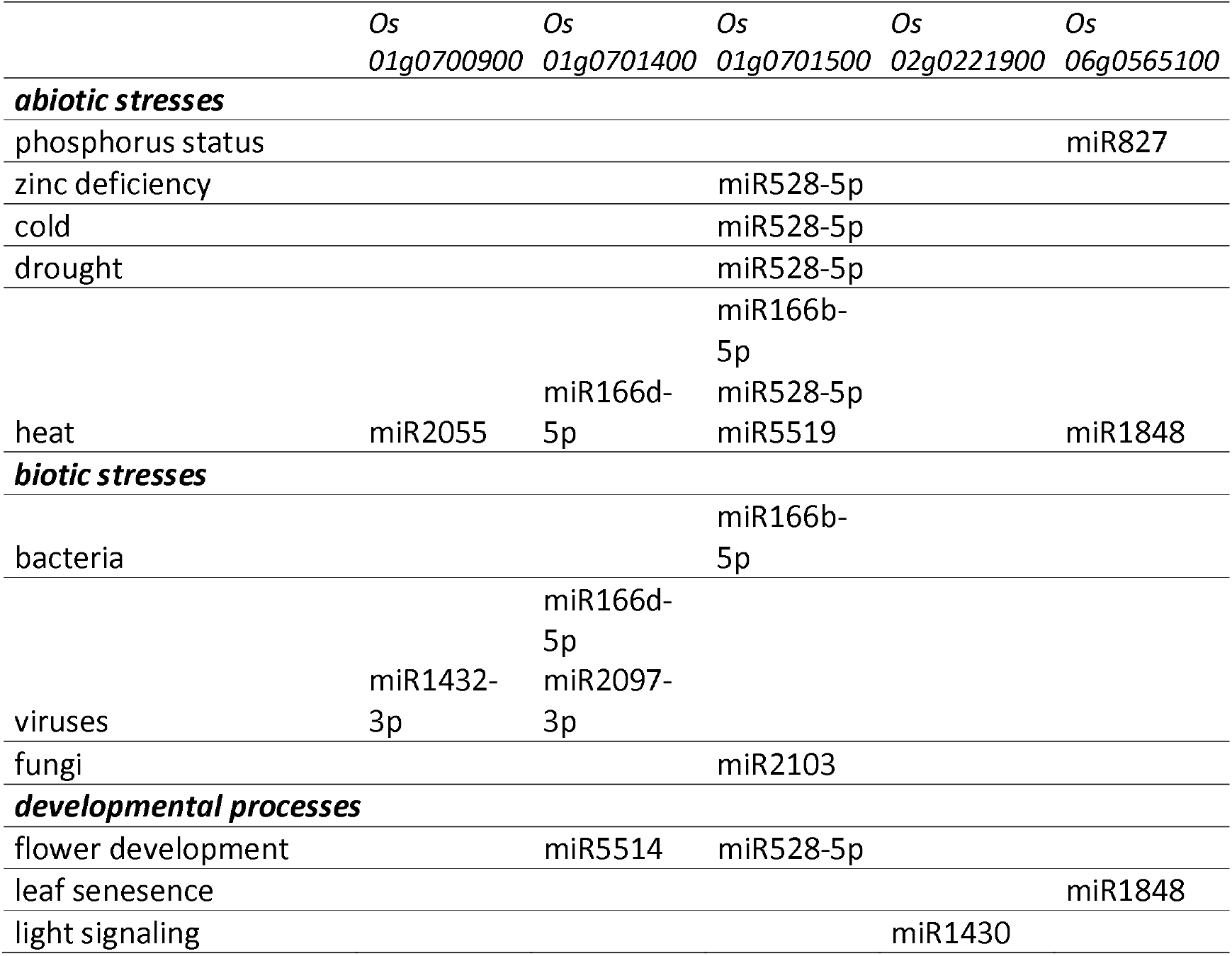
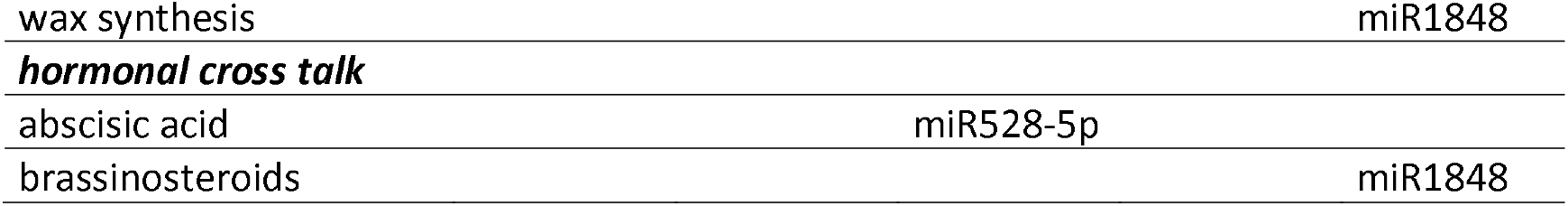
MiRNAs that bind *MAX1* homologues categorized according to the functions that play in plant development, growth and response to stresses.

### 3.9. MiRNA specific to Os01g0701400

Only four miRNAs (within nine identified) were specific to *Os01g0701400* (Figure 5a). Within these miRNA’s one was involved in abiotic stress response [136], two in biotic stress response [137,138] and one in developmental process [139,140]. Among abiotic stresses, response to heat was described for osa-miR166d-5p (Table 2), however in contrast to the previously described miRNA specific to *Os01g0700900* (osa-miR2055), in that case expression of osa-miR166d-5p was inhibited during heat treatment in tolerant variety Nagina 22 [136]. On the other hand, expression of osa-miR166d-5p increased more than 36 fold, in response to rice stripe virus infection [137]. Also, when rice plants were infected by southern rice black-streaked dwarf virus, the expression of another miRNA increased, osa-miR2097-3p, which was specific only to this *MAX1* gene [138]. Those experimental data suggest that both miRNAs (osa-miR166d-5p and osa-miR2097-3p) are involved in plant resistance to viruses (Table 2). Finally, expression of osa-miR5514 was two-fold higher in pollen (during meiosis) of diploid rice, when compared with autotetraploid [140], this feature suggests the role of osa-miR5514 in pollen development (Table 2).

### 3.10. MiRNA specific to Os01g0701500

Within the eight miRNAs identified for *Os01g0701500*, six were specific to this *MAX1* gene. Within these miRNAs, two were associated with responses to biotic stresses: bacteria (osa-miR166b-5p) [141] and fungi (osa-miR2103) infections [142] (Table 2). A wide range of abiotic stresses were represented by identified miRNAs, including plant responses to heat: osa-miR166b-5p, osa-miR528-5p [136] and osa-miR5519 [143], drought: osa-miR528-5p [144], cold: osa-miR528-5p [145], and zinc deficiency: osa-miR528-5p [146]. Experimental data indicated that a single miRNA, osa-miR528-5p, was upregulated when rice plants were exposed to zinc starvation [146], but was also upregulated over four-fold by cold temperature stress in cold-tolerant rice variety (Hitomebore) [145], and additionally plays the role in drought response (Table 2). Osa-miR528-5p showed the opposite differential expression pattern between drought-tolerant (Vandana, Aday Sel) and drought-susceptible (IR64) rice varieties. In leaf tissues, it was down-regulated in drought-tolerant rice varieties but up-regulated in drought-susceptible rice variety. What is also important is that the expression level of osa-miR528-5p was significantly higher in drought-tolerant rice varieties compared to a drought-susceptible rice variety under control conditions [144]. The explanation of this wide range of stress responses in which miR528-5p is involved, might be the fact that this miRNA is ABA-responsive (Table 2). It was shown that in the tissue of rice ABA deficient mutant *Osaba1*, the expression of miR528-5p is significantly upregulated (almost five-fold) when compared to the wild-type plants [147]. Finally, osa-miR528-5p was characterized as involved in flower development in rice [140] (Table 2). Studies carried out on di- and autotetraploid rice revealed that osa-miR528-5p plays a role in the early stage of pollen development through silencing the expression of the target genes (i.e., *LOC_Os06g06050*), and therefore resulted in abnormal pollen development in autotetraploid rice [140].

### 3.11. MiRNA specific to Os02g0221900

Within miRNAs identified for *Os02g0221900*, five were specific to this one *MAX1* gene. Within these miRNAs one (osa-miR1430) was functionally characterized. It was proved that osa-miR1430 and osa-miR169 may target the same gene which encoding a nuclear transcription factor Y subunit (*LOC_Os12g42400*) [148]. Since members of osa-miR169 were upregulated in the *phyB* mutant [148], osa-miR1430 may also be involved in phytochromeB-mediated light signalling pathway (Table 2).

### 3.12. MiRNA specific for Os06g0565100

Within the seven miRNAs identified for *Os06g0565100* only four were specific for this one *MAX1* gene. Osa-miR827 is a well-known regulator of rice responses to phosphorus deficiency [149,150]. Under phosphorus starvation the accumulation of osa-miR827 was observed in both shoots and roots, however stronger expression was noted in shoots [149,150]. For the second miRNA that was specific only for *Os06g0565100*, four possible functions were proposed: involved in response to heat [136], wax synthesis [151], leaf senescence [152] and crosstalk with brassinosteroids [153] (Table 2). In the case of heat tolerance, expression of osa-miR1848 was observed only in the tissues of a heat-tolerance variety during exposition to the high temperature [136]. Analyses of expression patterns in leaves of two super hybrid rice, Nei-2-You 6 (N2Y6, age-resistant rice) and LiangYou-Pei 9 (LYP9, age-sensitive rice) revealed that osa-miR1848 is involved in leaf senescence via targeting NAC TFs [152]. On the other hand the known target of osa-miR1848 is *OsWS1* (*Oryza sativa wax synthase isoform 1*) encoding a protein involved in cuticular wax formation [151]. It was shown that in the leaves of osa-miR1848 overexpressing plants, *OsWS1* expression decreased approximately five-fold compared with wild type plants [151]. Finally, the obtusifoliol 14α-demethylase gene OsCYP51G3 was described as a target of osa-miR1848. *OsCYP51G3* is involved in phytosterol biosynthesis, which is a precursor of brassinosteroids. Thus, overexpression of osa-miR1848 reduced brassinosteoids biosynthesis, because the relative amounts of six brassinosteroids (teasterone, 3-dehydroteasterone, typhasterol, 6-deoxoteasterone, 6-deoxotyphasterol, and castasterone) decreased in plants overexpressing osa-miR1848, when compared with wild-type plants [153].

### 3.13. Profile expression of MAX1 homologues

Gene expression data of Nipponbare rice variety from 284 experiments, carried out on plants at different age (from 3-days-old seedling, up to flowering time), different organs (shoot, root, leaf, panicle, anther, callus, seeds), and different treatment (i.e., drought, phosphorus starvation or cadmium treatment) were analysed (Figure 6,Table S9). In the case of 181 experiments, expression data were available for all five rice *MAX1* genes. In the case of 28 experiments the expression of all five *MAX1* homologues were upregulated (i.e., in the panicle before flowering, in roots of 7-, 21- and 35-day-old seedlings exposed to phosphorus deficiency) and in the case of 7 experiments, expression of all five rice *MAX1* genes was downregulated (i.e., in untreated leaves of 14-day-old seedlings, in shoots of 14-day-old seedling exposed to phosphorus starvation) (Table S9). On the other hand, in 61% of the experiments (111 from 181) the expression of one *MAX1* gene was different in comparison to the other four MAX1 homologues (Table S9). When 10-day-old seedlings were exposed to cadmium (100 μM CdCl_2_), expression of *Os02g0221900* remained upregulated after 1h and 24h after treatment, whereas expression of four other *MAX1* genes remained downregulated (project ID DRP001141) (Table S9). On the other hand, in mature seeds, induced expression was observed only for *Os01g0701500*, whereas expression of others *MAX1s* was repressed (project ID SRP028376). Similar results were obtained for leaf tissue culture (Project ID SRP017256). Whereas in anthers, after and before flowering, only the expression of *Os02g0221900* was upregulated (project ID SRP047482 and DRP001762). Other experiments when the expression profile of the single *MAX1* was different in comparison to the other four *MAX1* homologues, specific time points of phosphorus starvation were carried out, in both roots and shoots of rice plants at different ages (Table S9).

**Figure 6.**
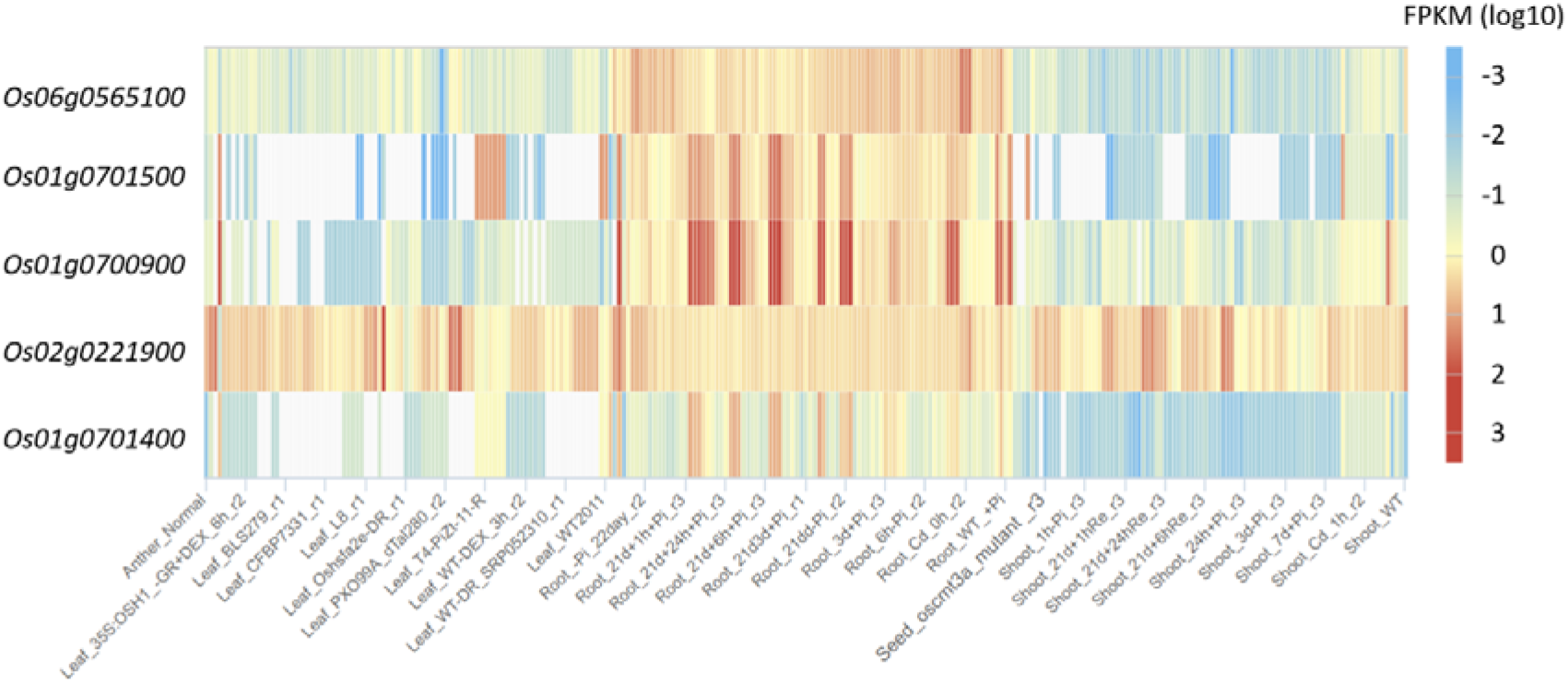
Heat map chart of rice *MAX1* gene expression profiles. FPKM - the number of fragments aligned per kilobases of the transcript per million mappable fragments from the total dataset. Description of the experiments was given in Table S9. White spots represent lack of data in this specific experiment to a given gene.

### 3.14. Co-expression gene networks of MAX1 homologues

For each of the *MAX1* homologues, a list of 27201 co-expressed genes was obtained (Table S10). The two *MAX1* homologues, *Os01g0700900* and *Os01g0701400*, demonstrated strong co-expression and were listed on the first positions of their co-expression gene network (CGN) list (Figure 7a, Table S10). These two are also phylogenetically closely related (Figure 3). No other *MAX1* homologs reach the criteria of MR lower then 100 (Figure 7a). In the case of four MAX1 homologs (*Os01g0700900, Os01g0701400, Os02g0221900, Os06g0565100*) more than 40 genes in CGN with MR lower than 100 were identified. Whereas in the case of *Os01g0701500* only 10 genes met these requirements (Figure 7b). Consistent with the high co-expression pattern of *Os01g0700900* and *Os01g0701400*, the highest number (14 of common genes was present in the CGNs of both *MAX1s*. No common genes from CGN with MR lower than 100 were found between *Os01g0701500*, *Os02g0221900* and *Os06g0565100* (Figure 7c).

**Figure 7.**
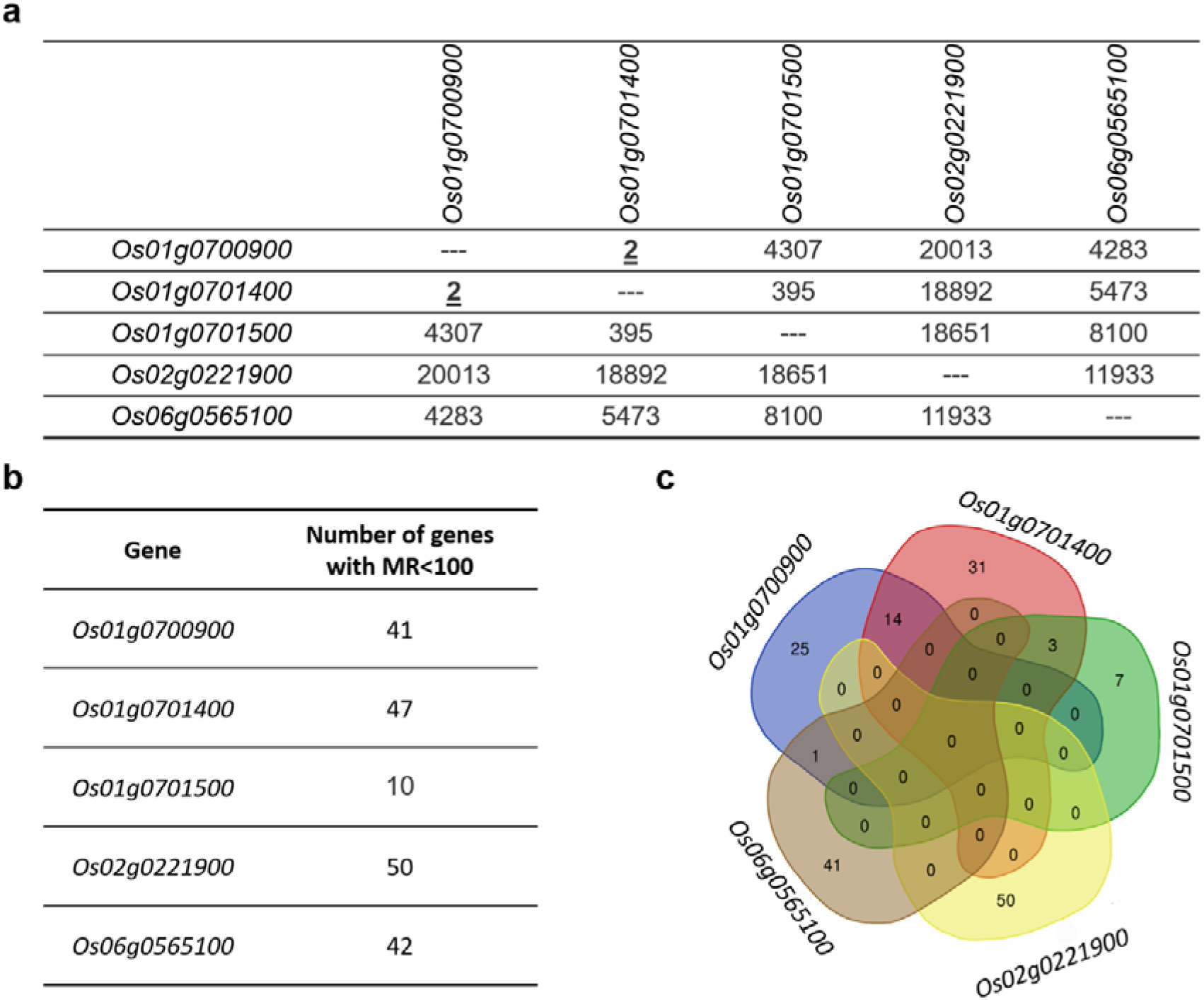
Co-expression gene network (CGN) of *MAX1* genes. (a) Mutual Rank (MR) of cooxpression between *MAX1s*. (b) Number of genes that exhibited similar expression pattern, such as *MAX1* homologues (genes with MR lower than 100). (c) The Venn diagram illustrates gens from CGN that are specific to each of rice *MAX1* genes and those that are common for different *MAX1* homologues.

### 3.15. Co-expression gene network of Os01g0700900

The analysis revealed that 25 genes were highly co-expressed (MR lower then 100) exclusively with *Os01g0700900*. Three genes from CGN of *Os01g0700900*, were described as involved in rice response to cadmium: *Os08g0106300* (MR=16.793, 2^nd^ position in list of co-expressed genes) [154], *Os08g0189200* (MR=91.913) [155] and *Os05g0162000* (MR= 94.106) [125] (Table 3). Moreover, gene ontology (GO) analysis also revealed the gene *Os04g0400800* (MR=16.793, 3^rd^ position in list of co-expressed genes) that encodes a protein that contains a heavy metal transport/detoxification domain (Table S10). Finally, *Os03g0149000* (MR=99.287), *Os06g0185500* (MR= 80.833) and *Os12g0454800* (MR= 85.229) play a role in rice response to ferulic acid [156], copper [157] and chromium [95], respectively. Changed expression of *Os05g0162000* that encodes a peroxidase, was observed not only in case of the rice response to cadmium, but also in the case of exposure to an excess of iron [129] and defence against rice blast fungus Magnaporthe oryzae [158] (Table 3). Another gene from *Os01g0700900* CGN: *Os03g0368900* (MR=85.229) encoding a peroxidase precursor is involved in ROS homeostasis in rice and is under the control of MADS3 (Os01g0201700) that regulates late anther development and pollen formation [159]. Finally, *Os10g0390800* (MR= 94.106), encoding another peroxidase (Table S10), plays the role in rice response to drought [52], phosphorus starvation [84] and bacterial infection [88] (Table 3).

**Table 3.**
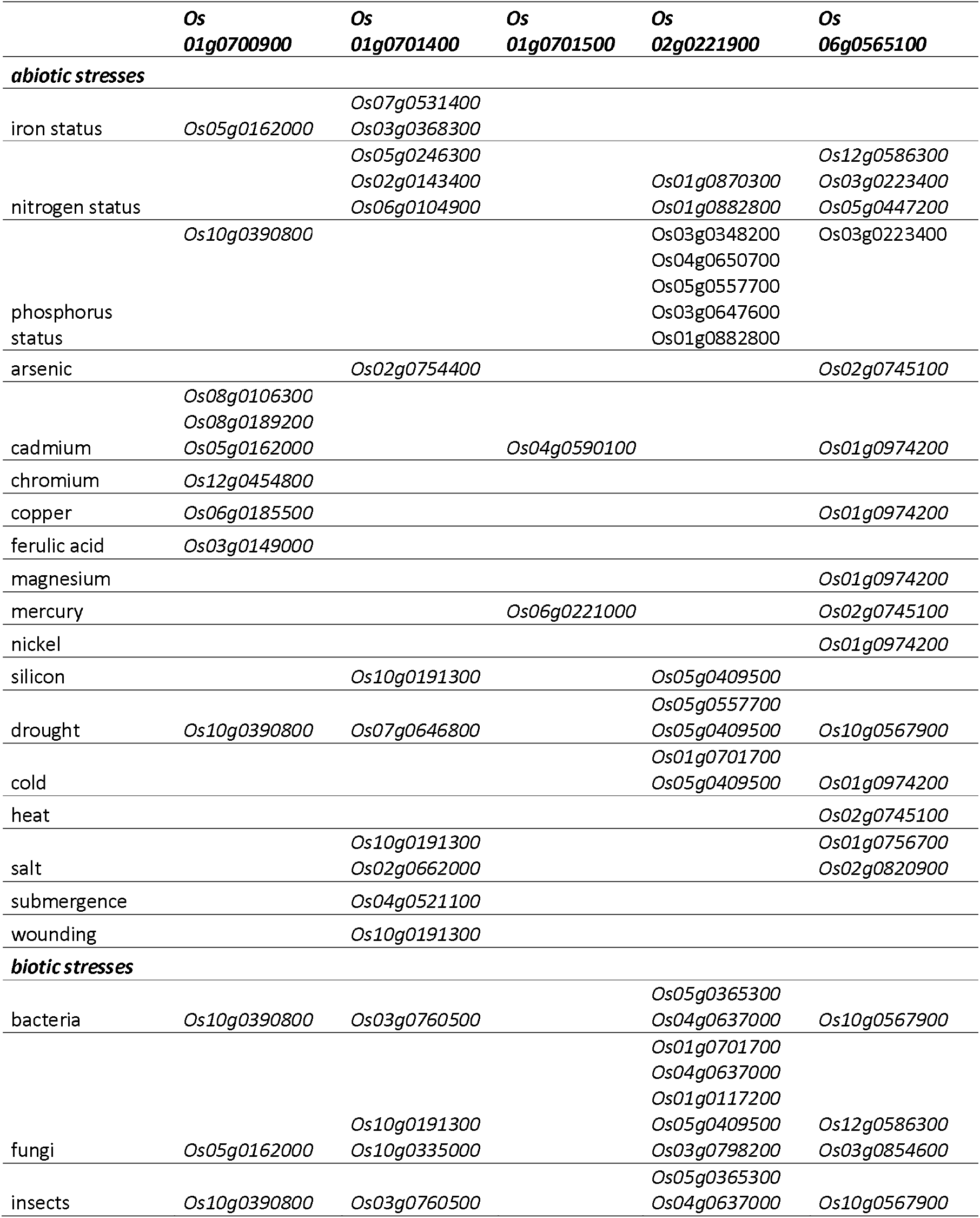
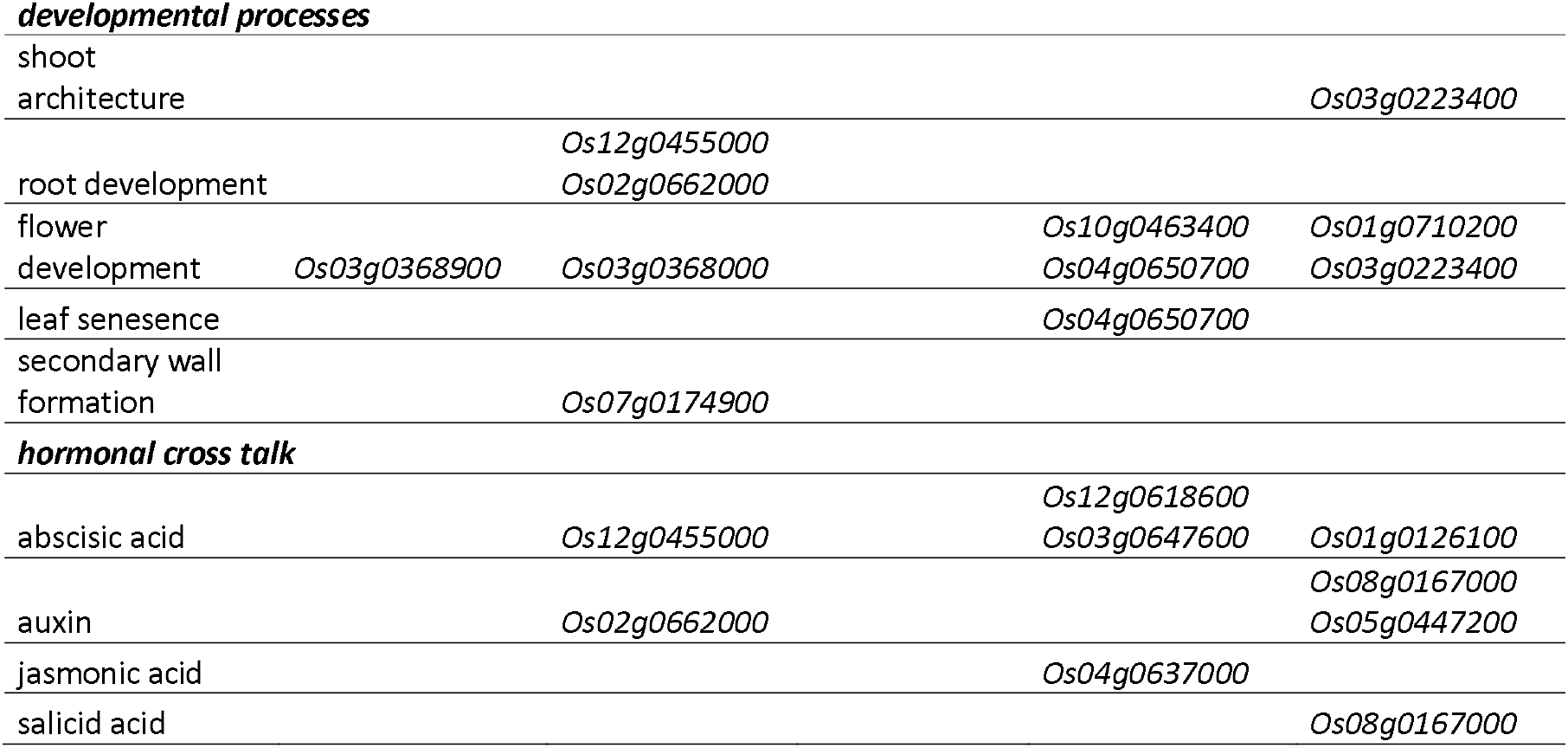
Function of genes co-expressed with *MAX1* homologues (MR<100).

### 3.16 Co-expression gene network of Os01g0701400

In the CGN of *Os01g0701400* (MR < 100) 31 genes were specific only to this *MAX1* homologue (Figure 7c). One of the identified genes, *Os07g0531400* (MR=72.595), was described in three independent experiments as a component of rice responses to deficiency or excess of iron [61,129,160], similarly to *Os03g0368300* (MR=39.987) [61] (Table 3). Also, genes involved in response to nitrogen status, *Os05g0246300* (MR=75.736) [161], *Os02g0143400* (MR=82.432) [60] and *Os06g0104900* (MR=75.736) [162] were found in the CGN of *Os01g0701400* (Table 3). Two other genes, *Os02g0754400* (MR=95.812) and *Os10g0191300* (MR= 81.425) play a role in response to arsenic [79] and silicon [163], respectively (Table 3). However, it was postulated that, *Os10g0191300*, which encodes a pathogenesis related protein (Table S10), might be involved in rice defence against insects [164] and fungi [163]. Other pathogen resistance genes, are, *Os03g0760500* (MR=96.156) - against bacteria [116], and *Os10g0335000* (MR=81.994) – against fungi [165]. A group of genes related to rice response to abiotic stresses was found among CGN of Os01g0701400, including salinity (*Os10g0191300*, MR= 81.425) [166], submergence (*Os04g0521100*, MR=38.21) [167], drought (*Os07g0646800*, MR=77.421) [53], as well as wounding (*Os10g0191300*, MR= 81.994) [168] (Table 3).

Another gene, *Os07g0174900* (MR=81.994), was strongly expressed in the outer part of roots, and thus could be involved in suberin and lignin biosynthesis [169]. Whereas according to the UniProt Database three additional genes are related with cell wall formation, *Os05g0246300, Os10g0335000* and *Os03g0368000* (MR=20.881) (UniProt). *Os01g0701400* was expressed during anther development and pollen formation (*Os03g0368000*) [159]. Whereas high ABA-induced expression of *Os12g0455000* (MR=20.881) was observed during root hair elongation [170]. Finally, analysis of rice lines with overexpression of *Os02g0662000* (MR= 71.498), which encodes Rcc3, a proline–rich protein (PRP), revealed that Os02g0662000 promotes development of the root system via increased accumulation of auxin in the root, and additionally increased tolerance to salt stress [171].

### 3.17. Co-expression gene network of Os01g0701500

Seven (from 10) genes found in CGN of *Os01g0701500* that met the criteria MR < 100 were specific to this *MAX1* homolog. Among them, a function is described in literature for only two. The first one is: *Os06g0221000* (MR=33.764), which encodes one of the TF belongs to MYB family (MYB-81), and its expression was upregulated by mercury (25 μM) treatment [172], and the second one - *Os04g0590100* (MR=35.567) was involved in rice responses to cadmium [155]. The protein encoded by this gene contains a heavy metal transport/detoxification domain (Table S10).

### 3.18. Co-expression gene network of Os02g0221900

All 50 genes from CGN of *Os02g0221900* with MR<100 were specific only for this one *MAX1* homologue (Figure 7a). Among that number, experimental data of the possible functions were available for 14 genes (Table S10). The largest group of genes was involved in response to biotic stresses, including fungi (*Os01g0701700*, MR=41.952 [173], *Os04g0637000*, MR=15.1, [174], *Os01g0117200*, MR=57.35, [175]; *Os05g0409500*, MR=64.312 [176]; *Os03g0798200*, MR=90.111, [177]), bacteria (*Os05g0365300*, MR=89.778, [178], *Os04g0637000* [179]) and insects (*Os04g0637000* [174]) (Table S3, Table S10). A wide range of abiotic stresses was represented in CGN of *Os02g0221900*, including response to cold (*Os01g0701700* [123]; *Os05g0409500* [124]), drought (*Os05g0557700* [180], *Os05g0409500*, [53]), phosphorus status (*Os03g0348200*, MR=50.279, [181]; *Os04g0650700*, MR=20, [181]; *Os05g0557700* [182,183]; *Os03g0647600*, MR=74.162 [184], *Os01g0882800*, MR= 98.122 [60]), nitrogen status (*Os01g0870300*, MR=90.427, [185]; *Os01g0882800* [186]), and silicone (*Os05g0409500*, [163]) (Table 3, Table S10). On the other hand, *Os10g0463400* (MR=61.425), encoding a B-type response regulator (Edh1, Early headingdate1), is involved in rice flowering. It was shown that the loss of function of Edh1 led to prolongation of the vegetative growth [187]. Whereas *OsASNase2* (*Os04g0650700*), encoding asparaginase2, is involved in asparagine catabolism, and plays a role in development of rice grains. *OsASNase2* is the major asparaginase isoform in rice shoots, and is expressed in the dorsal vascular bundles and in developing grains, as well as in mesophyll and phloem companion cells of senescent flag leaves [188]. Finally, two genes (*Os12g0618600*, MR=67.149 and *Os03g0647600*), were described as targets of ABA-responsive miRNA in rice [147], whereas *Os04g0637000* encoding TF OsTGAP1, that is induced under jasmonic acid treatment, and plays a role in biosynthesis of diterpenoid phytoalexin [189].

### 3.19. Co-expression gene network of Os06g0565100

Among 42 genes from *Os06g0565100* CGN (MR<100), 41 were specific only to this *MAX1* homologue (Figure 7c), and for them, 14 functions were proposed, based on the available literature (Table 3, Table S10). Again, the group of genes related to abiotic stresses was the largest. Expression of the one of the genes (*Os01g0974200*, MR=63.718) encoding metallothionein (OsMT2c) was induced by cadmium, nickel, magnesium, copper, and cold [125,190], whereas *Os02g0745100* (MR=28.142) plays a role in response to arsenic [79,191,192], mercury [172], and heat [193] (Table 3). Also, genes involved in responses to phosphorus status (*Os03g0223400*, MR=46.174, [186]) and nitrogen status (*Os12g0586300*, MR= 44.125 [194]; *Os03g0223400* [162]; *Os05g0447200*, MR=51.827 [195]), drought (*Os10g0567900*, MR=96.343, [196]) and salt stress (*Os01g0756700*, MR=34.467 [197]; *Os02g0820900*, MR=31.177) (Table 3) were identified. Os01g0756700 that enhances salt tolerance in rice, is encoding Shaker family K^+^ channel KAT1 (OsKAT1). The expression of *OsKAT1* was restricted to internodes and rachides of wild-type rice, whereas other Shaker family genes were expressed in various organs [197]. Some of the genes from CGN of *Os06g0565100* were also involved in rice responses to pathogen attack, such as bacteria (*Os10g0567900* [178]), insects (*Os01g0974200* [198]) and fungi (*Os12g0586300* [63], *Os03g0854600*, MR=33.764 [199]). Finally, genes related to developmental processes were identified in CGN of analysed *MAX1*. *Os01g0710200* (MR=88.978) that encodes polyamine oxidase7 (OsPAO7) is specifically expressed in anthers, with an expressional peak at the bicellular pollen stage [200]. On the other hand, *Os03g0223400* (MR=46.174), encoding glutamine synthetase (OsGS1;2), is involved in the development of rice grains [188,201] and glutamine-dependent tiller outgrowth [202]. Another gene, *Os01g0126100* (MR=21.977), was described as a target of ABA-responsive miRNA [147]. Whereas expression of *Os08g0167000* (MR=13.711), which encodes one of the half-size adenosine triphosphate-binding cassette transporter subgroup G (ABCG), was induced by salicid acid and repressed by auxin treatment [203]. Finally, *Os05g0447200* is *AUX2*, one of auxin-responsive genes, and was highly induced under microgravity conditions [204].

## 4. Discussion

The number of developmental processes and abiotic/biotic stresses that are SL-dependent are rapidly being uncovered [13]. The majority of our knowledge about the role of SL in plants comes from the analysis of mutants disturbed in SL biosynthesis and signalling. In the case of the biosynthesis mutants, plants with disorders in core biosynthesis pathway are usually used, because inactivation of the enzymes involved in initial stages of SL production results with lack/decreased amount of all SLs in plants [33]. However, each plant produces different SLs, and the composition of produced SL mixture may vary due to developmental stage or environmental conditions [205]. Moreover new SLs are still being discovered [15], and the question why plants are producing different classes of SLs is still open. Up to now, the origins of structural diversity of SLs remains unknown. The core of SL biosynthesis (up to CL) is highly conserved in mono- and dicots, whereas the next stages that result in the formation of various SLs, are different [23]. In *A. thaliana* MAX1 catalyses the oxidation of CL to CLA [19], which is next converted into an unknown SL-like product [33]. On the other hand, in rice there are five *MAX1* homologues, and it was postulated that four of them might be involved in different steps of SL biosynthesis [35]. However, the functions of the different produced SLs have not been fully reported so far, and specific enzymes involved in their biosynthesis have not been described yet.

Previously we have analysed rice and *A. thaliana* genes encoding proteins involved in SL biosynthesis, which allowed us to propose the new functions of SLs, and to describe mechanisms that may regulate their expression [11]. So far, various research reports that confirmed our predictions were published (reviewed by [12,206]). Here, we focused on all five rice *MAX1* genes. Our *in silico* analyses revealed the set of TFs and miRNAs that may be involved in the regulation of *MAX1* homologues in rice. Finally, analysis of RNA-seq data was used to describe the profiles of *MAX1* expression and reveal the genes from their co-expression gene networks. All the data indicate that there are specific mechanisms that regulate the expression of each single *MAX1*, and additionally, processes specific for individual MAX1s can be proposed.

### 4.1. TFs that regulate expression of MAX1 homologues

Because TFs are able to regulate the spatial expression of the many different genes, TFs play a crucial role in the coordination of plant growth, development and plant responses to different stresses [207]. More than 200 TFs that recognize motifs in the promoters of single *MAX1* were identified (Figure 4). Among them almost 25% (60 motifs) were common for all *MAX1s*, which indicates that in many cases expression of all five homologues exhibits the same regulation mechanisms (Figure 4b). Despite this, it was possible to select TF binding sites for each of the *MAX1* homologues that were specific only to a single *MAX1* gene. It should be noted that binding sites might be recognized by different TFs that belongs to the same family (Tables S3-S6). Thus, the expression of each of the *MAX1* homologues could be induced or repressed under specific regulatory mechanisms. Analysis of the TFs that were specific to single *MAX1s* revealed that some of them were already functionally characterized (Tables S3-S6) (Table 1). For example, one of the *MAX1* homologues (*Os01g0701500*) is under control of two TFs that act as master regulators of secondary wall formation in rice: SECONDARY WALL NAC DOMAIN PROTEIN1 and 2 (OsSWN1, Os01g0701500 and OsSWN2, Os06g0131700). It was shown that *OsSWN1* and *OsSWN2* are expressed in cells where secondary cell walls are formed and can alter secondary cell wall formation in rice: *OsSWN1* is highly active in sclerenchymatous cells of the leaf blade and less active in xylem cells, whereas OsSWN2 is highly active in xylem cells and less active in sclerenchymatous cells [107]. Additionally, two other TFs (OsSWN3, Os08g0103900 and OsSWN7, Os06g0104200) that are involved in cell wall biosynthesis, also bind this *MAX1* exclusively. Zhong and co-workers demonstrated that overexpression of rice *OsSWN1, OsSWN3* and *OsSWN7* genes in *A. thaliana* induces ectopic deposition of cellulose, xylan and lignin in secondary walls. These data indicated that the abovementioned three TFs are the main players in secondary wall formation in rice [108] and all of them regulate expression of single *MAX1* gene (Table 3). Thus, it might be speculated that this *MAX1* homolog (*Os01g0701500*) is involved in secondary wall formation in rice. On the other hand, another *MAX1* homologue (*Os02g0221900*) is under regulation of one of the major determinant of the seed specificity - OsVP1 (Os01g0911700). Analysis of temporal and spatial expression patterns of the *OsVP1*, revealed that activity of this TF could be detected in embryos as early as 2-3 days after pollination (DAP) and thereafter became preferentially localized to shoot, radicle and vascular tissues during the embryo development at both the mRNA and protein levels. Whereas in the aleurone layers, *OsVP1* mRNA and protein were detected after 6 DAP [114]. It was also shown that *OsVP1* is required for the induction of ABA-regulated genes that include genes encoding the late embryogenesis abundant (lea) protein [208]. Based on these results the role of another single *MAX1* homologue (*Os02g0221900*) during rice seed formation and development can be postulated. Both described functions, secondary wall formation and seed development were not previously reported for SLs. This might be because there are ‘non-canonical’ SL functions – the functions in which specific SLs, produced by a single MAX1, are involved, and the respective single mutants have not yet been analysed.

It also should be noted that TFs that exclusively bind the promoter region of a single *MAX1* may be also involved in the same processes as the TFs identified for other *MAX1* homologues. However, the presence of the unique TFs motifs in the promoter region of different *MAX1s* indicates that, under specific conditions, their expression will be regulated by different mechanisms.

### 4.2. Rice MAX1s are regulated by different miRNAs

MicroRNAs (miRNAs) are short, non-coding RNAs that identify complementary sites in mRNAs and target selected mRNAs for repression [209]. In contrast to the TFs, miRNAs were unique for each of the *MAX1*. For each of *MAX1*, at least eight miRNAs were identified, and what is more important, at least four of them were specific only to the mRNA of single *MAX1* (Figure 5). Moreover, none of the identified miRNAs were able to bind four or all five rice *MAX1* genes. Only two miRNAs recognize motifs in three *MAX1* homologues: osa-miR419 (*Os01g0700900, Os01g0701400, Os01g0701500*) and osa-miR5075 (*Os01g0701500, Os02g0221900, Os06g0565100*) (Table S8). Based on the known function of the miRNAs that may regulate *MAX1*, processes can be postulated that may involve specific *MAX1s*. For example, the mRNA of *Os01g0701500* is matched by osa-miR528-5p, which is involved in wide range of developmental processes and stress responses (Table 2). Analysis of the rice ABA deficient mutant, *Osaba1*, revealed a five-fold increase in expression of osa-miR528-5p compared to the wild-type [147]. Thus, an ABA-dependent miRNA may regulate the expression of gene encoding enzyme involved in SL biosynthesis. Recently, Visentin et al., described that exogenous application of SL results in the accumulation of miR156 in tomato, and moreover, an increase in guard cell sensitivity to ABA and stomatal closure was observed [210]. Here, we find out that an ABA-dependent miRNA may regulate one of the SL biosynthesis genes. This feature may indicate further crosstalk between SLs and ABA.

On the other hand, a second *MAX1* homologue - *Os06g0565100* – may be under the regulation of osa-miR1848 (Table 2). This miRNA plays a role in various processes including wax biosynthesis. This observation is in agreement with results obtained for TFs that regulate expression of *Os06g0565100*, including *OsWR1*, which promotes the expression of genes involved in wax synthesis. There is increasing evidence that SLs are involved in plant response to drought ([210–212] and one of the components of plant responses to drought is deposition of waxes. Thus, it can be proposed that one of the MAX1 homologue plays a role in wax biosynthesis and deposition, which may help plants to adapt to drought conditions.

### 4.3. Expression profiles of MAX1s and their co-expression gene networks

In general, the expression profiles of the analysed *MAX1* homologues were similar, however it should be noted that all five *MAX1* genes were not analysed in all the RNA-seq datasets (Table S9). As was expected, the induced expression of *MAX1* genes was observed when rice plants were exposed to nitrogen deficiency. However, this induction was mainly observed in roots for four *MAX1* homologues: *Os01g0700900*, *Os01g0701500*, *Os02g0221900* and *Os06g0565100*. Whereas, under the same conditions, the expression of *Os01g0701400* in root tissues decreased (Table S9). On the other hand, some of experimental data regarding gene expression confirmed our *in silico* prediction regarding specific roles of single *MAX1* genes. For example, only *Os02g0221900* expression was upregulated in seeds, whereas the transcription of all other *MAX1s* was downregulated (Table S9). Previously we showed that *Os02g0221900* is under regulation of the main seed-specific TF (*OsVP1*). RNA-seq data confirmed that only this *MAX1* is highly active during seed development. This strengthens our postulate that the regulation of seed development could be a non-canonical SL function in plants, which is mediated by *Os02g0221900*.

Analysis of genes that are co-expressed with individual *MAX1s* revealed that each *MAX1* homologue belongs to its own CGN that contains a number of genes specific to that *MAX1*. Interestingly, only in the CGN of *Os01g0700900* and Os01g0701500 (when MR<100) the other *MAX1* (*Os01g0701500* and *Os01g0700900*) was found (Figure 7). The data confirmed that *MAX1* genes are likely controlled by different regulatory mechanisms. Investigation of the function of genes that are co-expressed with *MAX1s* suggest that *Os01g0700900*, for example, could be involved in rice responses to various heavy metals (such as cadmium, chromium, copper), whereas other *MAX1s* are co-expressed with genes involved in responses to arsenic (*Os01g0701400*), cadmium and mercury (*Os01g0701500*), or arsenic, cadmium, copper, magnesium, mercury and nickel (*Os06g0565100*). Thus, it can be speculated that SLs play a role in plant responses to heavy metals that were not previously reported. Incidentally, the Bala rice cultivar (deletion of *Os01g0700900* and *Os01g0701400* on chromosome 1) expresses susceptibility to germanium toxicity [213]. Four of the five *MAX1s* were also co-expressed with genes that are related to pathogen (bacteria, fungi, insects) attack. That role of SLs was previously proposed [11] and some experimental data in this field were also published (reviewed by [12]). On the other hand, in the CGNs of four *MAX1* homologs, genes involved in flower development were also found, including, for example, a regulator of flowering (*Edh1, Early headingdate1*) and an enzyme specific for anthers and developing pollens (*OsPAO7, polyamine oxidase7*). Thus, new canonical functions of SLs that relate to flower development or responses to heavy metals can be postulated.

## 5. Conclusions

Based on *in silico* analysis and the analysis of RNA-seq experiments we suggest that the five *MAX1* homologues could be involved in different developmental processes and stress responses in rice. The predicted ‘non-SL-canonical’ functions in plants can now be confirmed by analysis of mutants in single *MAX1* genes. MAX1s are involved in the late stages of SL biosynthesis and they are responsible for much of the structural diversity of SLs. Thus, the prediction is that the products of MAX1 activity will play a role in different process. Here, we describe mechanisms that may regulate transcription and post-transcriptional regulation of each of the *MAX1* homologues in rice. The obtained data indicate the presence of regulatory mechanisms that are specific to single *MAX1s*, and also highlight candidates (TFs and miRNA) for further investigation about the role of single *MAX1* genes. Finally, analysis of *MAX1* expression profiles and genes that are co-expressed with *MAX1* homologues in rice provide some experimental support of our *in silico* predictions. Based on those data, new ‘SL-canonical’ functions in rice can be postulated, such as the regulation of flower development or responses to heavy metals. On the other hand, the presented data allows us to also suggest some of ‘non-SL-canonical’ functions related to single *MAX1* homologues (products of the enzymatic activity of single *MAX1s*), such as wax biosynthesis and the regulation of seed development.

Do the individual SL products of MAX1s and other SL biosynthesis enzymes act on specific plant traits? Our uncovering of the divergent nature of the potential pathways that could be regulating the individual MAX1s may indicate functional divergence, rather than simply redundancy or escape from parasitic weeds. CL and CLA are mobile and non-bioactive. Specific enzymes, like MAX1, may convert CLA to specific bioactive SLs near the site of action, which may induce tissue-specific responses. This is reminiscent of the gibberellin (GA) pathway where there are multiple different chemical forms, and major transported or stored forms, such as GA_12_, GA_20_ and GA_53_, are non-bioactive, and are converted to bioactive forms, such as GA_1_ and GA_4_, on-site [214,215]. Moreover, in bud outgrowth, GA_1_ and GA_4_ can promote bud elongation, whereas GA_3_ and GA_6_ seem to inhibit bud growth by deactivating GA_1_ and GA_4_ [216]. Other plant hormones also show diversity of chemical structure and activity. For example, isopentenyl-adenine and *trans*-zeatin cytokinins appear to have different functions [217].

Our predictions of new functions for SLs, and descriptions of the regulatory mechanisms for each of *MAX1* homologues, will facilitate the design of experiments, particularly using single mutants (in *MAX1s*, or TFs and miRNAs that are specific to the single *MAX1*). Also, analysis of the levels of different SLs that are produced and/or exuded by different genotypes needs to be performed during development processes or during responses to various treatments. Natural variation provided by the Bala rice cultivar could be a good place to begin those experimental approaches in combination with tissue-specific complementation.

## Supplementary Materials

[…]

## Acknowledgements

Thanks to Kaori Yoneyama and Koichi Yoneyama for discussions.

## Author Contributions

M.M., A.S., P.B.B. and A.B-Z. analysed the data and wrote the manuscript.

## References

1 Cook, C.E. et al. (1966) Germination of Witchweed (Striga lutea Lour.): Isolation and Properties of a Potent Stimulant. Science 154, 1189–1190

2 Akiyama, K. et al. (2005) Plant sesquiterpenes induce hyphal branching in arbuscular mycorrhizal fungi. Nature 435, 824–827

3 Gomez-Roldan, V. et al. (2008) Strigolactone inhibition of shoot branching. Nature 455, 189–194

4 Umehara, M. et al. (2008) Inhibition of shoot branching by new terpenoid plant hormones. Nature 455, 195–200

5 Sun, H. et al. (2016) The role of strigolactones in root development. Plant Signal Behav 11, e1110662

6 Hu, Z. et al. (2010) Strigolactones negatively regulate mesocotyl elongation in rice during germination and growth in darkness. Plant Cell Physiol. 51, 1136–1142

7 de Saint Germain, A. et al. (2013) Strigolactones stimulate internode elongation independently of gibberellins. Plant Physiol. 163, 1012–1025

8 Agusti, J. et al. (2011) Strigolactone signaling is required for auxin-dependent stimulation of secondary growth in plants. Proc. Natl. Acad. Sci. U.S.A. 108, 20242–20247

9 Zhang, J. et al. (2020) Strigolactones inhibit auxin feedback on PIN-dependent auxin transport canalization. Nat Commun 11, 3508

10 Yamada, Y. and Umehara, M. (2015) Possible Roles of Strigolactones during Leaf Senescence. Plants (Basel) 4, 664–677

11 Marzec, M. and Muszynska, A. (2015) In silico analysis of the genes encoding proteins that are involved in the biosynthesis of the RMS/MAX/D pathway revealed new roles of Strigolactones in plants. Int J Mol Sci 16, 6757–6782

12 Marzec, M. (2016) Strigolactones as Part of the Plant Defence System. Trends Plant Sci. 21, 900–903

13 Mostofa, M.G. et al. (2018) Strigolactones in plant adaptation to abiotic stresses: An emerging avenue of plant research. Plant Cell Environ. 41, 2227–2243

14 Borghi, L. et al. (2016) The importance of strigolactone transport regulation for symbiotic signaling and shoot branching. Planta 243, 1351–1360

15 Yoneyama, K. et al. (2018) Which are the major players, canonical or non-canonical strigolactones? J. Exp. Bot. 69, 2231–2239

16 Al-Babili, S. and Bouwmeester, H.J. (2015) Strigolactones, a novel carotenoid-derived plant hormone. Annu Rev Plant Biol 66, 161–186

17 Yoneyama, K. et al. (2020) Hydroxyl carlactone derivatives are predominant strigolactones in Arabidopsis. Plant Direct 4, e00219

18 Alder, A. et al. (2012) The path from β-carotene to carlactone, a strigolactone-like plant hormone. Science 335, 1348–1351

19 Abe, S. et al. (2014) Carlactone is converted to carlactonoic acid by MAX1 in Arabidopsis and its methyl ester can directly interact with AtD14 in vitro. Proc. Natl. Acad. Sci. U.S.A. 111, 18084–18089

20 Charnikhova, T.V. et al. (2017) Zealactones. Novel natural strigolactones from maize. Phytochemistry 137, 123–131

21 Xie, X. et al. (2017) Methyl zealactonoate, a novel germination stimulant for root parasitic weeds produced by maize. J Pestic Sci 42, 58–61

22 Kim, H.I. et al. (2014) Avenaol, a germination stimulant for root parasitic plants from Avena strigosa. Phytochemistry 103, 85–88

23 Wang, Y. and Bouwmeester, H.J. (2018) Structural diversity in the strigolactones. J. Exp. Bot. 69, 2219–2230

24 Yoneyama, K. et al. (2009) Strigolactones: structures and biological activities. Pest Manag. Sci. 65, 467–470

25 Jamil, M. et al. (2011) Quantification of the relationship between strigolactones and Striga hermonthica infection in rice under varying levels of nitrogen and phosphorus: N and P effect on strigolactones and Striga hermonthica. Weed Research 51, 373–385

26 Xie, X. et al. (2013) Confirming stereochemical structures of strigolactones produced by rice and tobacco. Mol Plant 6, 153–163

27 Boyer, F.-D. et al. (2012) Structure-activity relationship studies of strigolactone-related molecules for branching inhibition in garden pea: molecule design for shoot branching. Plant Physiol. 159, 1524–1544

28 Butt, H. et al. (2018) Engineering plant architecture via CRISPR/Cas9-mediated alteration of strigolactone biosynthesis. BMC Plant Biol. 18, 174

29 Marzec, M. (2016) Perception and Signaling of Strigolactones. Front Plant Sci 7, 1260

30 Marzec, M. and Brewer, P. (2019) Binding or Hydrolysis? How Does the Strigolactone Receptor Work? Trends Plant Sci. 24, 571–574

31 Booker, J. et al. (2005) MAX1 encodes a cytochrome P450 family member that acts downstream of MAX3/4 to produce a carotenoid-derived branch-inhibiting hormone. Dev. Cell 8, 443–449

32 Simons, J.L. et al. (2007) Analysis of the DECREASED APICAL DOMINANCE genes of petunia in the control of axillary branching. Plant Physiol. 143, 697–706

33 Brewer, P.B. et al. (2016) LATERAL BRANCHING OXIDOREDUCTASE acts in the final stages of strigolactone biosynthesis in Arabidopsis. Proc. Natl. Acad. Sci. U.S.A. 113, 6301–6306

34 Iseki, M. et al. (2018) Evidence for species-dependent biosynthetic pathways for converting carlactone to strigolactones in plants. J. Exp. Bot. 69, 2305–2318

35 Yoneyama, K. et al. (2018) Conversion of carlactone to carlactonoic acid is a conserved function of MAX1 homologs in strigolactone biosynthesis. New Phytol. 218, 1522–1533

36 Zhang, Y. et al. (2018) The tomato MAX1 homolog, SlMAX1, is involved in the biosynthesis of tomato strigolactones from carlactone. New Phytol. 219, 297–309

37 Zheng, M. et al. (2020) Knockout of two BnaMAX1 homologs by CRISPR/Cas9-targeted mutagenesis improves plant architecture and increases yield in rapeseed (Brassica napus L.). Plant Biotechnol. J. 18, 644–654

38 Mori, N. et al. (2020) Chemical identification of 18-hydroxycarlactonoic acid as an LjMAX1 product and in planta conversion of its methyl ester to canonical and non-canonical strigolactones in Lotus japonicus. Phytochemistry 174, 112349

39 Zhang, Y. et al. (2014) Rice cytochrome P450 MAX1 homologs catalyze distinct steps in strigolactone biosynthesis. Nat. Chem. Biol. 10, 1028–1033

40 Cardoso, C. et al. (2014) Natural variation of rice strigolactone biosynthesis is associated with the deletion of two MAX1 orthologs. Proc. Natl. Acad. Sci. U.S.A. 111, 2379–2384

41 Wakabayashi, T. et al. (2019) Direct conversion of carlactonoic acid to orobanchol by cytochrome P450 CYP722C in strigolactone biosynthesis. Sci Adv 5, eaax9067

42 Wakabayashi, T. et al. (2020) CYP722C from Gossypium arboreum catalyzes the conversion of carlactonoic acid to 5-deoxystrigol. Planta 251, 97

43 Challis, R.J. et al. (2013) A role for more axillary growth1 (MAX1) in evolutionary diversity in strigolactone signaling upstream of MAX2. Plant Physiol. 161, 1885–1902

44 Walker, C.H. et al. (2019) Strigolactone synthesis is ancestral in land plants, but canonical strigolactone signalling is a flowering plant innovation. BMC Biol. 17, 70

45 Katoh, K. et al. (2002) MAFFT: a novel method for rapid multiple sequence alignment based on fast Fourier transform. Nucleic Acids Res. 30, 3059–3066

46 Dai, X. et al. (2018) psRNATarget: a plant small RNA target analysis server (2017 release). Nucleic Acids Research 46, W49–W54

47 da Maia, L.C. et al. (2017) Transcriptome profiling of rice seedlings under cold stress. Functional Plant Biol. 44, 419

48 Wang, H. et al. (2018) Genome-wide association study reveals candidate genes related to low temperature tolerance in rice (Oryza sativa) during germination. 3 Biotech 8, 235

49 Kitazumi, A. et al. (2018) Potential of Oryza officinalis to augment the cold tolerance genetic mechanisms of Oryza sativa by network complementation. Sci Rep 8, 16346

50 Shin, S.-J. et al. (2016) Novel drought-responsive regulatory coding and non-coding transcripts from Oryza Sativa L. Genes Genom 38, 949–960

51 Ahn, H. et al. (2017) Transcriptional Network Analysis Reveals Drought Resistance Mechanisms of AP2/ERF Transgenic Rice. Front Plant Sci 8, 1044

52 Wang, D. et al. (2011) Genome-wide temporal-spatial gene expression profiling of drought responsiveness in rice. BMC Genomics 12, 149

53 Chung, P.J. et al. (2018) Genome-wide analyses of direct target genes of four rice NAC-domain transcription factors involved in drought tolerance. BMC Genomics 19, 40

54 Kim, S.-W. et al. (2015) Integrating omics analysis of salt stress-responsive genes in rice. Genes Genom 37, 645–655

55 Mito, T. et al. (2011) Generation of chimeric repressors that confer salt tolerance in Arabidopsis and rice. Plant Biotechnol. J. 9, 736–746

56 Hossain, M.Rashed. (2014), Salinity tolerance and transcriptomics in rice., University of Birmingham

57 Das, C.K. et al. (2018) Computational analysis of genes encoding for molecular determinants of arsenic tolerance in rice (*Oryza sativa* L.) to engineer low arsenic content varieties. ORYZA-An Internatio. Journ. on Rice 55, 248

58 Ogawa, I. et al. (2009) Time course analysis of gene regulation under cadmium stress in rice. Plant Soil 325, 97–108

59 Yang, H.-C. et al. (2017) Identification of early ammonium nitrate-responsive genes in rice roots. Sci Rep 7, 16885

60 Hsieh, P.-H. et al. (2018) Early molecular events associated with nitrogen deficiency in rice seedling roots. Sci Rep 8, 12207

61 Bashir, K. et al. (2014) Transcriptomic analysis of rice in response to iron deficiency and excess. Rice (N Y) 7, 18

62 Kikuchi, S. (2014) Genome-wide view of the expression profiles of NAC-domain genes in response to infection by rice viruses. In Omics technologies and crop improvement (Benkeblia, N., ed), pp. 127–152, CRC Press

63 Wang, C. et al. (2019) Transcriptome analysis of a rice cultivar reveals the differentially expressed genes in response to wild and mutant strains of Xanthomonas oryzae pv. oryzae. Sci Rep 9, 3757

64 Tezuka, D. et al. (2019) The rice ethylene response factor OsERF83 positively regulates disease resistance to Magnaporthe oryzae. Plant Physiol. Biochem. 135, 263–271

65 Lavarenne, J. et al. (2019) Inference of the gene regulatory network acting downstream of CROWN ROOTLESS 1 in rice reveals a regulatory cascade linking genes involved in auxin signaling, crown root initiation, and root meristem specification and maintenance. Plant J. 100, 954–968

66 Matsubara, K. and Yano, M. (2018) Genetic and Molecular Dissection of Flowering Time Control in Rice. In Rice Genomics, Genetics and Breeding (Sasaki, T. and Ashikari, M., eds), pp. 177–190, Springer Singapore

67 Das, S.P. et al. (2018) Micromorphic and Molecular Studies of Floral Organs of a Multiple Seeded Rice (Oryza sativa L.). Plant Mol Biol Rep 36, 764–775

68 Lee, S. et al. (2017) Molecular bases for differential aging programs between flag and second leaves during grain-filling in rice. Sci Rep 7, 8792

69 Yamatani, H. et al. (2013) NYC4, the rice ortholog of Arabidopsis THF1, is involved in the degradation of chlorophyll - protein complexes during leaf senescence. Plant J. 74, 652–662

70 Sugimoto, K. et al. (2009) Genetic Control of Seed Dormancy in Rice. In Gamma Field Symposia

71 Chu, Y. et al. (2019) Rice transcription factor OsMADS57 regulates plant height by modulating gibberellin catabolism. Rice (N Y) 12, 38

72 Nguyen, T.D. et al. (2016) Genome-wide identification and analysis of rice genes preferentially expressed in pollen at an early developmental stage. Plant Mol. Biol. 92, 71–88

73 Kubo, T. et al. (2013) Transcriptome analysis of developing ovules in rice isolated by laser microdissection. Plant Cell Physiol. 54, 750–765

74 Fang, J. et al. (2019) Ef-cd locus shortens rice maturity duration without yield penalty. Proc. Natl. Acad. Sci. U.S.A. 116, 18717–18722

75 Tsuji, H. et al. (2011) Regulation of flowering in rice: two florigen genes, a complex gene network, and natural variation. Curr. Opin. Plant Biol. 14, 45–52

76 Hori, K. et al. (2016) Genetic control of flowering time in rice: integration of Mendelian genetics and genomics. Theor. Appl. Genet. 129, 2241–2252

77 Kitomi, Y. et al. (2018) Genetic Mechanisms Involved in the Formation of Root System Architecture. In Rice Genomics, Genetics and Breeding (Sasaki, T. and Ashikari, M., eds), pp. 241–274, Springer Singapore

78 Lee, D.-K. et al. (2016) Overexpression of the OsERF71 Transcription Factor Alters Rice Root Structure and Drought Resistance. Plant Physiol. 172, 575–588

79 Singh, P.K. et al. (2017) Nitric oxide mediated transcriptional modulation enhances plant adaptive responses to arsenic stress. Sci Rep 7, 3592

80 Zhang, F. et al. (2012) Genome-wide gene expression profiling of introgressed indica rice alleles associated with seedling cold tolerance improvement in a japonica rice background. BMC Genomics 13, 461

81 Tula, S. et al. (2013) Physiological Assessment and Allele Mining in Rice Cultivars for Salinity and Drought Stress Tolerance. Vegetos-An Inter. Jour. of Plnt. Rese. 26, 219

82 Mohanty, B. et al. (2016) Identification of candidate network hubs involved in metabolic adjustments of rice under drought stress by integrating transcriptome data and genome-scale metabolic network. Plant Sci. 242, 224–239

83 Xu, Y. et al. (2017) Transcriptional regulation of hormone-synthesis and signaling pathways by overexpressing cytokinin-synthesis contributes to improved drought tolerance in creeping bentgrass. Physiol Plant 161, 235–256

84 Oono, Y. et al. (2013) Diversity in the complexity of phosphate starvation transcriptomes among rice cultivars based on RNA-Seq profiles. Plant Mol. Biol. 83, 523–537

85 Mohanty, B. et al. (2016) Transcriptional regulatory mechanism of alcohol dehydrogenase 1-deficient mutant of rice for cell survival under complete submergence. Rice (N Y) 9, 51

86 Croissant-Sych, Y. and Okita, T.W. (1996) Identification of positive and negative regulatory cis-elements of the rice glutelin Gt3 promoter. Plant Science 116, 27–35

87 Mohanty, B. et al. (2005) Detection and preliminary analysis of motifs in promoters of anaerobically induced genes of different plant species. Ann. Bot. 96, 669–681

88 King, E. et al. (2019) Monitoring of Rice Transcriptional Responses to Contrasted Colonizing Patterns of Phytobeneficial Burkholderia s.l. Reveals a Temporal Shift in JA Systemic Response. Front Plant Sci 10, 1141

89 Le, C.T.T. et al. (2016) ZINC FINGER OF ARABIDOPSIS THALIANA12 (ZAT12) Interacts with FER-LIKE IRON DEFICIENCY-INDUCED TRANSCRIPTION FACTOR (FIT) Linking Iron Deficiency and Oxidative Stress Responses. Plant Physiol. 170, 540–557

90 Kan, C.-C. et al. (2015) Glutamine rapidly induces the expression of key transcription factor genes involved in nitrogen and stress responses in rice roots. BMC Genomics 16, 731

91 Fujino, K. and Matsuda, Y. (2010) Genome-wide analysis of genes targeted by qLTG3-1 controlling low-temperature germinability in rice. Plant Mol. Biol. 72, 137–152

92 Cui, Y. et al. (2018) OsDSSR1, a novel small peptide, enhances drought tolerance in transgenic rice. Plant Sci. 270, 85–96

93 Gao, W. et al. (2019) Gene Expression Profiles Deciphering the Pathways of Coronatine Alleviating Water Stress in Rice (Oryza sativa L.) Cultivar Nipponbare (Japonica). Int J Mol Sci 20,

94 Tuteja, N. et al. (2015) OsBAT1 Augments Salinity Stress Tolerance by Enhancing Detoxification of ROS and Expression of Stress-Responsive Genes in Transgenic Rice. Plant Mol Biol Rep 33, 1192–1209

95 Huang, T.-L. et al. (2014) Genomic profiling of rice roots with short- and long-term chromium stress. Plant Mol. Biol. 86, 157–170

96 Xiong, H. et al. (2012) Comparative transcriptional profiling of two rice genotypes carrying SUB1A-1 but exhibiting differential tolerance to submergence. Functional Plant Biol. 39, 449

97 Yasui, Y. et al. (2017) Genetic Enhancer Analysis Reveals that FLORAL ORGAN NUMBER2 and OsMADS3 Co-operatively Regulate Maintenance and Determinacy of the Flower Meristem in Rice. Plant Cell Physiol. 58, 893–903

98 Ohnishi, T. et al. (2011) Distinct gene expression profiles in egg and synergid cells of rice as revealed by cell type-specific microarrays. Plant Physiol. 155, 881–891

99 Sato, H. et al. (2012) Male-sterile and cleistogamous phenotypes in tall fescue induced by chimeric repressors of SUPERWOMAN1 and OsMADS58. Plant Sci. 183, 183–189

100 Kuwano, M. et al. (2011) A novel endosperm transfer cell-containing region-specific gene and its promoter in rice. Plant Mol. Biol. 76, 47–56

101 Ke, S. et al. (2018) Genome-wide transcriptome profiling provides insights into panicle development of rice (*Oryza sativa* L.). Gene 675, 285–300

102 Kim, J.S. et al. (2017) Genome-wide identification of grain filling genes regulated by the OsSMF1 transcription factor in rice. Rice (N Y) 10, 16

103 Piao, W. et al. (2015) Rice Phytochrome B (OsPhyB) Negatively Regulates Dark- and Starvation-Induced Leaf Senescence. Plants (Basel) 4, 644–663

104 Lee, S.-H. et al. (2015) Mutation of Oryza sativa CORONATINE INSENSITIVE 1b (OsCOI1b) delays leaf senescence. J Integr Plant Biol 57, 562–576

105 Hirano, K. et al. (2013) Survey of Genes Involved in Rice Secondary Cell Wall Formation Through a Co-Expression Network. Plant and Cell Physiology 54, 1803–1821

106 Hirano, K. et al. (2013) Identification of Transcription Factors Involved in Rice Secondary Cell Wall Formation. Plant and Cell Physiology 54, 1791–1802

107 Yoshida, K. et al. (2013) Engineering the Oryza sativa cell wall with rice NAC transcription factors regulating secondary wall formation. Front. Plant Sci. 4,

108 Zhong, R. et al. (2011) Transcriptional activation of secondary wall biosynthesis by rice and maize NAC and MYB transcription factors. Plant Cell Physiol. 52, 1856–1871

109 Yang, C. et al. (2013) Overexpression of microRNA319 impacts leaf morphogenesis and leads to enhanced cold tolerance in rice (Oryza sativa L.). Plant Cell Environ. 36, 2207–2218

110 Jin, Y. et al. (2018) OsERF101, an ERF family transcription factor, regulates drought stress response in reproductive tissues. Plant Mol. Biol. 98, 51–65

111 Kortz, A. et al. (2019) Cell Type-Specific Transcriptomics of Lateral Root Formation and Plasticity. Front Plant Sci 10, 21

112 Gupta, M.K. et al. (2017) The impact of natural selection on gene associated with panicle number formation in Oryza sativa. Can J Biotech 1, 198–198

113 Fu, Z. et al. (2014) The Rice Basic Helix-Loop-Helix Transcription Factor TDR INTERACTING PROTEIN2 Is a Central Switch in Early Anther Development. Plant Cell 26, 1512–1524

114 Miyoshi, K. et al. (2002) Temporal and spatial expression pattern of the OSVP1 and OSEM genes during seed development in rice. Plant Cell Physiol. 43, 307–313

115 Minakuchi, K. et al. (2010) FINE CULM1 (FC1) works downstream of strigolactones to inhibit the outgrowth of axillary buds in rice. Plant Cell Physiol. 51, 1127–1135

116 Xu, Mei Rong, X. et al. (2011) Different patterns of gene expression in rice varieties undergoing a resistant or susceptible interaction with the bacterial leaf streak pathogen. Afr. J. Biotechnol. 10, 14419–14438

117 Brusamarello-Santos, L.C.C. et al. (2012) Differential gene expression of rice roots inoculated with the diazotroph Herbaspirillum seropedicae. Plant Soil 356, 113–125

118 Yi, S.Y. et al. (2013) Microarray Analysis of bacterial blight resistance 1 mutant rice infected with Xanthomonas oryzae pv. oryzae. Plant Breed. Biotech. 1, 354–365

119 Lilly, J. and Subramanian, B. (2019) Gene network mediated by WRKY13 to regulate resistance against sheath infecting fungi in rice (Oryza sativa L.). Plant Sci. 280, 269–282

120 He, H. et al. (2016) Comparative mapping of powdery mildew resistance gene Pm21 and functional characterization of resistance-related genes in wheat. Theor. Appl. Genet. 129, 819–829

121 Wang, Y. et al. (2012) Identification of transcription factors potential related to brown planthopper resistance in rice via microarray expression profiling. BMC Genomics 13, 687

122 Yuexiong, Z. et al. (2020) Identification of Major Locus Bph35 Resistance to Brown Planthopper in Rice. Rice Science 27, 237–245

123 Zhao, J. et al. (2015) Global transcriptional profiling of a cold-tolerant rice variety under moderate cold stress reveals different cold stress response mechanisms. Physiol Plant 154, 381–394

124 Suzuki, K. et al. (2015) Cooling water before panicle initiation increases chilling-induced male sterility and disables chilling-induced expression of genes encoding OsFKBP65 and heat shock proteins in rice spikelets. Plant Cell Environ. 38, 1255–1274

125 Sun, L. et al. (2019) Transcriptome analysis of rice (Oryza sativa L.) shoots responsive to cadmium stress. Sci Rep 9, 10177

126 Lakshmanan, M. et al. (2014) Metabolic and transcriptional regulatory mechanisms underlying the anoxic adaptation of rice coleoptile. AoB PLANTS 6,

127 Kottapalli, K.R. et al. (2007) Combining in silico mapping and arraying: an approach to identifying common candidate genes for submergence tolerance and resistance to bacterial leaf blight in rice. Mol. Cells 24, 394–408

128 Wu, J. et al. (2020) Overexpression of MADS-box transcription factor OsMADS25 enhances salt stress tolerance in Rice and Arabidopsis. Plant Growth Regul 90, 163–171

129 Finatto, T. et al. (2015) Abiotic stress and genome dynamics: specific genes and transposable elements response to iron excess in rice. Rice (N Y) 8, 13

130 Sinha, S.K. et al. (2018) Transcriptome Analysis of Two Rice Varieties Contrasting for Nitrogen Use Efficiency under Chronic N Starvation Reveals Differences in Chloroplast and Starch Metabolism-Related Genes. Genes (Basel) 9,

131 Fujishiro, Y. et al. (2018) Comprehensive panicle phenotyping reveals that qSrn7/FZP influences higher-order branching. Sci Rep 8, 12511

132 Jung, H. et al. (2017) Overexpression of OsERF48 causes regulation of OsCML16, a calmodulin-like protein gene that enhances root growth and drought tolerance. Plant Biotechnol. J. 15, 1295–1308

133 Lu, Z. et al. (2013) Genome-wide binding analysis of the transcription activator ideal plant architecture1 reveals a complex network regulating rice plant architecture. Plant Cell 25, 3743–3759

134 Wang, Y. et al. (2012) An ethylene response factor OsWR1 responsive to drought stress transcriptionally activates wax synthesis related genes and increases wax production in rice. Plant Mol Biol 78, 275–288

135 Yaish, M.W. et al. (2010) The APETALA-2-like transcription factor OsAP2-39 controls key interactions between abscisic acid and gibberellin in rice. PLoS Genet. 6, e1001098

136 Mangrauthia, S.K. et al. (2017) Genome-wide changes in microRNA expression during short and prolonged heat stress and recovery in contrasting rice cultivars. J. Exp. Bot. 68, 2399–2412

137 Yang, J. et al. (2016) Integrative Analysis of the microRNAome and Transcriptome Illuminates the Response of Susceptible Rice Plants to Rice Stripe Virus. PLoS ONE 11, e0146946

138 Xu, D. et al. (2014) MicroRNAs responding to southern rice black-streaked dwarf virus infection and their target genes associated with symptom development in rice. Virus Res. 190, 60–68

139 Ta, K.N. et al. (2016) miR2118-triggered phased siRNAs are differentially expressed during the panicle development of wild and domesticated African rice species. Rice (N Y) 9, 10

140 Li, X. et al. (2016) Comparative Small RNA Analysis of Pollen Development in Autotetraploid and Diploid Rice. Int J Mol Sci 17, 499

141 Li, W. et al. (2019) Integration Analysis of Small RNA and Degradome Sequencing Reveals MicroRNAs Responsive to Dickeya zeae in Resistant Rice. Int J Mol Sci 20,

142 Fan, J. et al. (2020) circRNAs Are Involved in the Rice-Magnaporthe oryzae Interaction. Plant Physiol. 182, 272–286

143 Li, J. et al. (2015) Genome-Wide Identification of MicroRNAs Responsive to High Temperature in Rice (*Oryza sativa*) by High-Throughput Deep Sequencing. J Agro Crop Sci 201, 379–388

144 Cheah, B.H. et al. (2015) Identification of four functionally important microRNA families with contrasting differential expression profiles between drought-tolerant and susceptible rice leaf at vegetative stage. BMC Genomics 16, 692

145 Maeda, S. et al. (2016) Comparative analysis of microRNA profiles of rice anthers between cool-sensitive and cool-tolerant cultivars under cool-temperature stress. Genes Genet. Syst. 91, 97–109

146 Zeng, H. et al. (2019) Integrated analyses of miRNAome and transcriptome reveal zinc deficiency responses in rice seedlings. BMC Plant Biol. 19, 585

147 Tian, C. et al. (2015) Identification and Characterization of ABA-Responsive MicroRNAs in Rice. J Genet Genomics 42, 393–402

148 Sun, W. et al. (2015) Genome-wide identification of microRNAs and their targets in wild type and phyB mutant provides a key link between microRNAs and the phyB-mediated light signaling pathway in rice. Front Plant Sci 6, 372

149 Lin, S.-I. et al. (2010) Complex regulation of two target genes encoding SPX-MFS proteins by rice miR827 in response to phosphate starvation. Plant Cell Physiol. 51, 2119–2131

150 Wang, C. et al. (2012) Functional characterization of the rice SPX-MFS family reveals a key role of OsSPX-MFS1 in controlling phosphate homeostasis in leaves. New Phytol. 196, 139–148

151 Xia, K. et al. (2015) OsWS1 involved in cuticular wax biosynthesis is regulated by osa-miR1848. Plant Cell Environ. 38, 2662–2673

152 Xu, X. et al. (2014) Genome-wide analysis of microRNAs and their target genes related to leaf senescence of rice. PLoS ONE 9, e114313

153 Xia, K. et al. (2015) Rice microRNA osa-miR1848 targets the obtusifoliol 14α-demethylase gene OsCYP51G3 and mediates the biosynthesis of phytosterols and brassinosteroids during development and in response to stress. New Phytol. 208, 790–802

154 Zhang, M. et al. (2012) Transcriptional profiling in cadmium-treated rice seedling roots using suppressive subtractive hybridization. Plant Physiol. Biochem. 50, 79–86

155 Lee, S.-B. et al. (2019) Functional Haplotype and eQTL Analyses of Genes Affecting Cadmium Content in Cultivated Rice. Rice (N Y) 12, 84

156 Chi, W.-C. et al. (2013) Autotoxicity mechanism of Oryza sativa: transcriptome response in rice roots exposed to ferulic acid. BMC Genomics 14, 351

157 Liu, Q. et al. (2015) Transcriptional and physiological analyses identify a regulatory role for hydrogen peroxide in the lignin biosynthesis of copper-stressed rice roots. Plant Soil 387, 323–336

158 Chujo, T. et al. (2013) OsWRKY28, a PAMP-responsive transrepressor, negatively regulates innate immune responses in rice against rice blast fungus. Plant Mol. Biol. 82, 23–37

159 Hu, L. et al. (2011) Rice MADS3 Regulates ROS Homeostasis during Late Anther Development. Plant Cell 23, 515–533

160 Aung, M.S. et al. (2018) Physiological and transcriptomic analysis of responses to different levels of iron excess stress in various rice tissues. Soil Science and Plant Nutrition 64, 370–385

161 Hayashi, T. et al. (2009) Effects of High Nitrogen Supply on the Susceptibility to Coolness at the Young Microspore Stage in Rice (*Oryza sativa* L.): Gene Expression Analysis in Mature Anthers. Plant Production Science 12, 271–277

162 Yang, S. et al. (2015) RNA-Seq analysis of differentially expressed genes in rice under varied nitrogen supplies. Gene 555, 305–317

163 Brunings, A.M. et al. (2009) Differential gene expression of rice in response to silicon and rice blast fungus Magnaporthe oryzae. Annals of Applied Biology 155, 161–170

164 Sun, X. et al. (2019) Transcriptome analysis of roots from resistant and susceptible rice varieties infected with Hirschmanniella mucronata. FEBS Open Bio 9, 1968–1982

165 Singh, V. et al. (2016) Comparative transcriptomics of rice and exploitation of target genes for blast infection. Agri Gene 1, 143–150

166 Lakra, N. et al. (2019) Mapping the ‘early salinity response’ triggered proteome adaptation in contrasting rice genotypes using iTRAQ approach. Rice 12, 3

167 Phule, A.S. et al. (2019) RNA-seq reveals the involvement of key genes for aerobic adaptation in rice. Sci Rep 9, 5235

168 Bertini, L. et al. (2019) Proteomic Analysis of MeJa-Induced Defense Responses in Rice against Wounding. Int J Mol Sci 20,

169 Shiono, K. et al. (2014) Microarray analysis of laser-microdissected tissues indicates the biosynthesis of suberin in the outer part of roots during formation of a barrier to radial oxygen loss in rice (Oryza sativa). J. Exp. Bot. 65, 4795–4806

170 Wang, T. et al. (2017) Abscisic Acid Regulates Auxin Homeostasis in Rice Root Tips to Promote Root Hair Elongation. Front Plant Sci 8, 1121

171 Li, X. et al. (2018) Overexpression of RCc3 improves root system architecture and enhances salt tolerance in rice. Plant Physiol. Biochem. 130, 566–576

172 Chen, Y.-A. et al. (2014) Transcriptome profiling and physiological studies reveal a major role for aromatic amino acids in mercury stress tolerance in rice seedlings. PLoS ONE 9, e95163

173 Tian, L. et al. (2019) Comparative study of the mycorrhizal root transcriptomes of wild and cultivated rice in response to the pathogen Magnaporthe oryzae. Rice (N Y) 12, 35

174 Verbeek, R.E.M. et al. (2019) Jasmonate-Induced Defense Mechanisms in the Belowground Antagonistic Interaction Between Pythium arrhenomanes and Meloidogyne graminicola in Rice. Front Plant Sci 10, 1515

175 Yokotani, N. et al. (2014) OsNAC111, a blast disease-responsive transcription factor in rice, positively regulates the expression of defense-related genes. Mol. Plant Microbe Interact. 27, 1027–1034

176 Tian, L. et al. (2018) Comparative analysis of the root transcriptomes of cultivated and wild rice varieties in response to Magnaporthe oryzae infection revealed both common and species-specific pathogen responses. Rice (N Y) 11, 26

177 Al-Bader, N. et al. (2019) Loss of premature stop codon in the Wall-Associated Kinase 91 (OsWAK91) gene confers sheath blight disease resistance in rice, Plant Biology.

178 Wang, X. et al. (2013) Characterization of a Novel NBS-LRR Gene Involved in Bacterial Blight Resistance in Rice. Plant Mol Biol Rep 31, 649–656

179 Li, W.Q. et al. (2018) Genome-wide identification and characterization of long non-coding RNAs responsive to Dickeya zeae in rice. RSC Adv. 8, 34408–34417

180 Chung, P.J. et al. (2016) Transcriptome profiling of drought responsive noncoding RNAs and their target genes in rice. BMC Genomics 17, 563

181 Han, J. et al. (2018) A mitochondrial phosphate transporter, McPht gene, confers an acclimation regulation of the transgenic rice to phosphorus deficiency. Journal of Integrative Agriculture 17, 1932–1945

182 Jeong, K. et al. (2017) Phosphorus remobilization from rice flag leaves during grain filling: an RNA-seq study. Plant Biotechnol. J. 15, 15–26

183 Ying, Y. et al. (2017) Two h-Type Thioredoxins Interact with the E2 Ubiquitin Conjugase PHO2 to Fine-Tune Phosphate Homeostasis in Rice. Plant Physiol. 173, 812–824

184 Takehisa, H. and Sato, Y. (2019) Transcriptome monitoring visualizes growth stage-dependent nutrient status dynamics in rice under field conditions. Plant J 97, 1048–1060

185 Obara, M. et al. (2011) Mapping quantitative trait loci controlling root length in rice seedlings grown with low or sufficient supply using backcross recombinant lines derived from a cross between *Oryza sativa* L. and *Oryza glaberrima* Steud. Soil Science and Plant Nutrition 57, 80–92

186 Shin, S.-Y. et al. (2018) Transcriptomic analyses of rice (Oryza sativa) genes and non-coding RNAs under nitrogen starvation using multiple omics technologies. BMC Genomics 19, 532

187 Wei, F.-J. et al. (2016) Both Hd1 and Ehd1 are important for artificial selection of flowering time in cultivated rice. Plant Sci. 242, 187–194

188 Yabuki, Y. et al. (2017) A temporal and spatial contribution of asparaginase to asparagine catabolism during development of rice grains. Rice (N Y) 10, 3

189 Yoshida, Y. et al. (2017) OsTGAP1 is responsible for JA-inducible diterpenoid phytoalexin biosynthesis in rice roots with biological impacts on allelopathic interaction. Physiol Plant 161, 532–544

190 Kim, Y.-O. and Kang, H. (2018) Comparative expression analysis of genes encoding metallothioneins in response to heavy metals and abiotic stresses in rice (Oryza sativa) and Arabidopsis thaliana. Biosci. Biotechnol. Biochem. 82, 1656–1665

191 Kuramata, M. et al. (2013) Genetic diversity of arsenic accumulation in rice and QTL analysis of methylated arsenic in rice grains. Rice (N Y) 6, 3

192 Huang, T.-L. et al. (2012) Transcriptomic changes and signalling pathways induced by arsenic stress in rice roots. Plant Mol. Biol. 80, 587–608

193 Zhang, X. et al. (2012) Expression profile in rice panicle: insights into heat response mechanism at reproductive stage. PLoS ONE 7, e49652

194 Sawaki, N. et al. (2013) A nitrate-inducible GARP family gene encodes an auto-repressible transcriptional repressor in rice. Plant Cell Physiol. 54, 506–517

195 Sun, L. et al. (2017) Spatio-temporal dynamics in global rice gene expression (Oryza sativa⍰ L.) in response to high ammonium stress. J. Plant Physiol. 212, 94–104

196 Redillas, M.C.F.R. et al. (2012) The overexpression of OsNAC9 alters the root architecture of rice plants enhancing drought resistance and grain yield under field conditions. Plant Biotechnol. J. 10, 792–805

197 Obata, T. et al. (2007) Rice shaker potassium channel OsKAT1 confers tolerance to salinity stress on yeast and rice cells. Plant Physiol. 144, 1978–1985

198 Wang, H.-L. et al. (2020) RNA-Seq revealed that infection with white tip nematodes could downregulate rice photosynthetic genes. Funct. Integr. Genomics 20, 367–381

199 Nasir, F. et al. (2019) Strigolactones positively regulate defense against Magnaporthe oryzae in rice (Oryza sativa). Plant Physiol. Biochem. 142, 106–116

200 Liu, T. et al. (2014) Polyamine oxidase 7 is a terminal catabolism-type enzyme in Oryza sativa and is specifically expressed in anthers. Plant Cell Physiol. 55, 1110–1122

201 You, C. et al. (2017) iTRAQ-based proteome profile analysis of superior and inferior Spikelets at early grain filling stage in japonica Rice. BMC Plant Biol. 17, 100

202 Ohashi, M. et al. (2018) Outgrowth of Rice Tillers Requires Availability of Glutamine in the Basal Portions of Shoots. Rice (N Y) 11, 31

203 Matsuda, S. et al. (2012) Genome-wide analysis and expression profiling of half-size ABC protein subgroup G in rice in response to abiotic stress and phytohormone treatments. Mol. Genet. Genomics 287, 819–835

204 Wakabayashi, K. et al. (2017) Persistence of plant hormone levels in rice shoots grown under microgravity conditions in space: its relationship to maintenance of shoot growth. Physiol Plantarum 161, 285–293

205 Ruyter-Spira, C. et al. (2013) The biology of strigolactones. Trends Plant Sci. 18, 72–83

206 Marzec, M. (2017) Strigolactones and Gibberellins: A New Couple in the Phytohormone World? Trends Plant Sci. 22, 813–815

207 Meshi, T. and Iwabuchi, M. (1995) Plant transcription factors. Plant Cell Physiol. 36, 1405–1420

208 Nakagawa, H. et al. (1996) The seed-specific transcription factor VP1 (OSVP1) is expressed in rice suspension-cultured cells. Plant Cell Physiol. 37, 355–362

209 Bartel, D.P. (2018) Metazoan MicroRNAs. Cell 173, 20–51

210 Visentin, I. et al. (2020) A novel strigolactone-miR156 module controls stomatal behaviour during drought recovery. Plant Cell Environ 43, 1613–1624

211 Haider, I. et al. (2018) The interaction of strigolactones with abscisic acid during the drought response in rice. J. Exp. Bot. 69, 2403–2414

212 Marzec, M. et al. (2020) Barley strigolactone signaling mutant hvd14.d reveals the role of strigolactones in ABA-dependent response to drought. Plant Cell Environ. DOI: 10.1111/pce.13815

213 Talukdar, P. et al. (2015) Biallelic and Genome Wide Association Mapping of Germanium Tolerant Loci in Rice (Oryza sativa L.). PLoS ONE 10, e0137577

214 Proebsting, W.M. et al. (1992) Gibberellin concentration and transport in genetic lines of pea⍰: effects of grafting. Plant Physiol. 100, 1354–1360

215 Regnault, T. et al. (2015) The gibberellin precursor GA12 acts as a long-distance growth signal in Arabidopsis. Nat Plants 1, 15073

216 Katyayini, N.U. et al. (2020) Dual Role of Gibberellin in Perennial Shoot Branching: Inhibition and Activation. Front Plant Sci 11, 736

217 Wheeldon, C.D. and Bennett, T. (2020) There and back again: An evolutionary perspective on long-distance coordination of plant growth and development. Semin. Cell Dev. Biol. DOI: 10.1016/j.semcdb.2020.06.011

